# The knowledge landscape of jaguar science: rapid growth, thematic structure, and regional inequalities

**DOI:** 10.64898/2026.08.17.745206

**Authors:** Marco Aurélio Mendes Elias, Philip Teles Soares, José Alexandre Felizola Diniz-Filho, Anah Teresa de Almeida Jácomo, Tiago Jácomo Silveira, Leandro Silveira

## Abstract

Understanding how scientific effort is distributed is essential for evaluating the evidence available to guide species conservation. We assessed the temporal, thematic, methodological, and spatial organization of jaguar (*Panthera onca*) research across the species’ range. Using 1,028 bibliographic records, we applied a metadata-based scope filter, defining a primary analytical corpus of 857 articles and an inclusive sensitivity corpus of 989. We analyzed publication growth from 1946–2024, author-keyword networks, high-specificity research-domain and methodological indicators, article-based spatial concentration across nine regions, and anthropogenic context using the Human Impact Index. Publication output increased strongly and nonlinearly, with the negative-binomial generalized additive model explaining 96% of deviance. Population and abundance, human–wildlife conflict and coexistence, and movement and connectivity were the most frequent research domains, whereas camera trapping was the most frequent methodological approach. Spatial analyses included 237 georeferenced terrestrial articles. Research was strongly concentrated in the Pantanal, Mesoamerica, and Atlantic Forest, but was underrepresented relative to range area in the Andes and Chocó–Darién, Amazon, and Guiana Shield. These patterns were stable to the broader scope definition. Research concentration showed no strong association with region-wide anthropogenic pressure, although studies within several extensive regions tended to occur in more human-influenced portions than the regional average. Jaguar research therefore shows substantial growth and partial alignment with applied conservation challenges but remains geographically uneven. Expanding representative research in underrepresented regions, while maintaining work in threatened landscapes, would strengthen the evidence base for range-wide conservation planning and improve coordination among countries, institutions, and regional research traditions.

## 1. INTRODUCTION

The increasing human transformation of terrestrial ecosystems has contributed to widespread declines of large carnivores, with consequences extending from population persistence to trophic interactions and ecosystem functioning (Estes *et al*., 2011; Ripple *et al*., 2014). The jaguar (*Panthera onca*), the largest felid in the Americas, has been extirpated from more than half of its historical range, with remaining populations increasingly affected by habitat conversion and fragmentation, prey depletion, hunting, persecution, and conflict with people (De La Torre *et al*., 2018; Romero-Muñoz *et al*., 2019). As an apex predator distributed across forests, wetlands, savannas, drylands, and increasingly human-modified landscapes, the jaguar has become a prominent focus of ecological research and conservation planning in the Neotropics. Its extensive range has also supported continental initiatives centered on the protection of core populations and the maintenance of landscape connectivity, with potential benefits extending to other species and ecosystems (Rabinowitz & Zeller, 2010; Thornton *et al*., 2016).

Research on jaguars has expanded substantially over recent decades. The adoption of systematic camera trapping, GPS telemetry, non-invasive genetics, remote sensing, and spatial modeling has increased the capacity to investigate abundance, movement, habitat use, connectivity, behavior, and interactions with people across diverse and often logistically challenging environments (Silver *et al*., 2004; Rabinowitz & Zeller, 2010; Roques *et al*., 2014; Morato *et al*., 2016). Together, these approaches expanded the spatial scale, temporal resolution, and range of ecological and conservation questions that could be investigated. However, growth in publication output alone does not ensure that knowledge is distributed evenly among research domains, methodological approaches, or geographic regions. Scientific effort may remain concentrated around particular questions, technologies, study systems, institutions, or locations, potentially leaving other components of a species’ ecology and distribution comparatively understudied (Wilson *et al*., 2016; Di Marco *et al*., 2017).

Evaluating the organization of this evidence base requires moving beyond publication counts alone. Quantitative scientometric approaches can describe the representation of research domains and methodological approaches, while author-keyword networks reveal how concepts are connected within the literature (Zupic & Čater, 2015; Aria & Cuccurullo, 2017). However, bibliometric prominence measures attention and connectivity within a defined corpus rather than the ecological importance, scientific quality, or conservation adequacy of a topic. A high frequency of studies addressing population status, movement, or human–wildlife conflict, for example, may reflect their direct relevance to management rather than an undesirable thematic imbalance. Conversely, limited representation alone does not demonstrate that a topic constitutes a priority knowledge gap. Because the patterns recovered by scientometric analyses also depend on corpus definition, metadata availability, and keyword standardization, transparent filtering and sensitivity analyses are essential for robust interpretation (Donthu *et al*., 2021). Scientometric evidence is therefore most informative when used to characterize the structure of the literature while keeping descriptive patterns distinct from normative judgments about future research priorities.

The geographic distribution of knowledge presents an additional challenge. Conservation research and biodiversity information are often concentrated in accessible and institutionally well-connected locations, with coverage also shaped by economic resources, language, geographic isolation, and security (Amano & Sutherland, 2013; Wilson *et al*., 2016). Within the Neotropics, for example, conservation research in the Caatinga has been shown to cluster near roads and research centers, illustrating how infrastructure can influence the geography of knowledge production (Lessa *et al*., 2019). For a species spanning much of the region, uneven research coverage may therefore reflect a combination of ecological, logistical, institutional, and conservation-related factors and should not automatically be interpreted as a failure to address conservation needs. Nevertheless, limited representation of extensive portions of the jaguar’s range may restrict the transferability of ecological inference and management recommendations across environmentally heterogeneous regions (Yates *et al*., 2018). Comparing observed article-based effort with the proportion of the mapped range contained within each region provides a transparent area-proportional expectation for identifying geographic concentration. This comparison identifies regions receiving more or less research attention than expected from their share of the jaguar’s range, without implying that research should be allocated strictly in proportion to area.

Anthropogenic pressure provides a second spatial benchmark for interpreting the geography of research because it allows us to contrast alternative mechanisms of knowledge production. Under a conservation-demand mechanism, scientific attention should increase where cumulative human modification generates more immediate ecological and management challenges, including habitat conversion, fragmentation, and interactions between people and wildlife. Under an opportunity-driven mechanism, however, research may concentrate where roads, settlements, institutions, and long-term field infrastructure make studies more feasible, irrespective of relative conservation need (Reddy & Dávalos, 2003; Kadmon *et al*., 2004; Wilson *et al*., 2016). These mechanisms are not mutually exclusive and may weaken a simple regional relationship between research concentration and anthropogenic pressure. Their expression may also be scale-dependent because regional averages can conceal the clustering of studies in more modified portions of otherwise extensive, comparatively low-impact regions, reflecting the broader dependence of ecological patterns on the spatial scale at which they are measured (Levin, 1992). Thus, it is important to examine both region-wide anthropogenic conditions and the conditions surrounding georeferenced study locations.

Here, we provide an integrative assessment of the temporal, thematic, methodological, and spatial organization of jaguar research. We combine publication trends, author-keyword frequencies and co-occurrence networks, metadata-derived indicators of research domains and methodological approaches, article-based spatial analyses across broad regions, and the Human Impact Index (Sanderson *et al*., 2002; Venter *et al*., 2016a, 2016b). We tested four hypotheses and their associated predictions. First, under the research-expansion hypothesis, we predicted that annual publication output would increase strongly and nonlinearly through time. Second, under the knowledge-structure hypothesis, we predicted that research would be unevenly distributed among domains and methods, with population and abundance, movement and connectivity, human–wildlife conflict and coexistence, and camera trapping among the most prominent components of the literature. We further predicted that author keywords would form identifiable conceptual communities rather than an unstructured network. Third, under the spatial-concentration hypothesis, we predicted that article-based research effort would depart from an area-proportional distribution across the jaguar’s range, resulting in regions with both greater and lower representation than expected from their geographic extent. Fourth, under the anthropogenic-alignment hypothesis, we predicted that research concentration would increase with region-wide anthropogenic pressure and that georeferenced studies would tend to occur in more human-influenced portions of each region. Together, these analyses evaluate whether the expansion of jaguar research has produced a thematically and geographically balanced evidence base and whether its spatial distribution corresponds to contemporary patterns of human influence.

## 2. METHODS

### 2.1. Literature search and analytical corpus

We assembled the source dataset through systematic searches of the Web of Science Core Collection (WoS; Clarivate) and Scopus (Elsevier), which provide broad and complementary coverage of peer-reviewed literature (Mongeon & Paul-Hus, 2016). Searches included records indexed through 23 June 2025 and used equivalent database-specific combinations of the scientific and common names “*Panthera onca*” OR “jaguar” OR “onça-pintada” in titles, abstracts, and keywords, without language restrictions. After merging records and removing duplicates, screening excluded non-article document types, records unrelated to the species, and studies without sufficient evidence that jaguars formed part of the ecological, conservation, or community-level analysis. Articles published in 2025 were excluded because that year was incomplete, yielding the 1946–2024 source dataset (**Figure S1**; **Table S1**).

Because records differed in their relevance to jaguar research, we applied a reproducible metadata-based scope filter. Titles, abstracts, author keywords, and indexed keywords were standardized and searched for multilingual jaguar terms. Records with jaguar terms in the title or author keywords were classified as high-confidence jaguar focus; those with at least two abstract mentions but no title or author-keyword evidence as comparative or multispecies jaguar focus; single abstract mentions as contextual or uncertain; and indexed-keyword-only or no-signal records as incidental. The primary analytical corpus combined the first two categories (n = 857), whereas the inclusive sensitivity corpus additionally retained contextual or uncertain records (n = 989); incidental records were excluded from both. Screening used titles and abstracts, with ambiguous records checked against full text or extended metadata. Operational definitions, corpus composition, and record-level classifications are provided in **Table S1**.

### 2.2. Research domains, methodological approaches, and author-keyword networks

We used two complementary approaches. First, predefined high-specificity metadata dictionaries independently classified nine non-exclusive research domains: population and abundance; movement and connectivity; human–wildlife conflict and coexistence; habitat loss, land-use change, and fragmentation; conservation planning and management; conservation genetics and genomics; health, disease, and pathogens; reproduction, ex situ conservation, and captive management; and diet and trophic ecology. Six methodological categories were also classified: camera trapping; telemetry and GPS tracking; genetic and molecular methods; spatial modelling and remote sensing; interviews and social-science methods; and capture–recapture estimation.

Primary classification required at least one corresponding expression in the title or author keywords. A broader classification used for sensitivity and auditing additionally accepted concordant evidence in both the abstract and indexed keywords. Articles could be assigned to multiple categories. High-specificity rules excluded generic taxonomic, phylogenetic, and pathogen-genomics mentions from conservation genetics unless accompanied by explicit population, landscape, diversity, connectivity, or conservation context. Complete dictionaries, classification rules, frequency summaries, and field-match audits are provided in **Table S2**.

Second, author keywords were used to characterize conceptual relationships within the literature. Keywords were standardized for capitalization, punctuation, hyphenation, and synonyms; search anchors, generic bibliographic or taxonomic terms, country names, and administrative geographic terms were excluded, while ecologically meaningful regional terms were retained. Repeated occurrences within an article were counted once. The 25 most frequent standardized keywords were summarized descriptively. For network analysis, we retained the 20 most frequent terms among those occurring in at least five articles. Unique within-article keyword pairs defined an undirected co-occurrence network weighted by Jaccard similarity; edges occurring in fewer than three articles were excluded. The graph two-core was retained when sufficiently large; otherwise, the largest connected component was used. Communities were identified with the Louvain algorithm, and node prominence was described by degree, weighted strength, and betweenness centrality (Blondel *et al*., 2008). Network analyses were implemented in igraph (Csardi & Nepusz, 2006). Full standardization rules, network metrics, community structure, and the author-plus-indexed-keyword sensitivity analysis are reported in **Table S3**.

### 2.3. Temporal analyses

Annual publication counts were calculated for the primary corpus from 1946 to 2024, including years with zero publications. We compared three models fitted to the same annual count response: negative-binomial intercept-only and linear models and a negative-binomial generalized additive model (GAM) with a smooth function of publication year. The intercept-only and linear negative-binomial models were fitted with MASS (Venables & Ripley, 2002), and the GAM was fitted by restricted maximum likelihood with mgcv (Wood, 2017). Models were compared by Akaike’s Information Criterion (AIC) and evaluated using effective degrees of freedom, deviance explained, adjusted R², and lag-one autocorrelation of deviance residuals. The selected model was used descriptively rather than to infer mechanisms of publication growth.

Temporal changes in research domains and methods were examined from 1990 onward to limit the influence of sparse early records. For each primary indicator, article-level presence or absence was modelled against publication year using a binomial GAM with a logit link. Models were fitted only when an indicator had at least 15 positive and 15 negative observations across at least ten publication years. Predicted annual probabilities and 95% confidence intervals were derived from fitted models, and observed proportions were grouped into five-year intervals for visualization. Smooth-term p-values were adjusted separately for research-domain and methodological families using the Benjamini–Hochberg procedure (Benjamini and Hochberg, 1995); these analyses were treated as exploratory. Scope sensitivity was evaluated by comparing annual publication counts between primary and inclusive corpora using Spearman’s rank correlation. Detailed specifications are provided in the Supplementary Methods; diagnostics, temporal results, annual counts, and sensitivity analyses are reported in **Table S4**.

### 2.4. Spatial data, regionalization, and research concentration

The coordinate database contained 1,762 rows associated with bibliographic records. Coordinates were converted to numeric form, invalid geographic values were excluded, and available manual-validation fields were incorporated. Retained coordinates were linked to bibliographic records using source identifiers, DOIs, or normalized article titles. Exact coordinate duplicates were retained for auditing but did not receive additional weight in article-based analyses; validation and linkage procedures are summarized in **Table S5**.

Coordinates were represented as WGS 84 points and intersected in R using sf (Pebesma, 2018) with the mapped jaguar distribution, terrestrial ecoregions, and country boundaries. Jaguar distribution followed the IUCN Red List map (Quigley *et al*., 2017; IUCN, 2026); source polygons were geometrically validated and dissolved into a single distribution geometry. Terrestrial ecoregions followed Olson *et al*. (2001) and were intersected with the jaguar range so that all area denominators referred only to mapped range; the generic “Lake” class was excluded. A reviewed crosswalk assigned ecoregions to 15 subregions and nine broad regions: Mesoamerica; Amazon; Guiana Shield; Andes and Chocó–Darién; Llanos and Orinoco; Atlantic Forest; Cerrado and Caatinga; Pantanal; and Chaco and Chiquitano dry forests. Areas were calculated in World Cylindrical Equal Area (EPSG:6933). The complete regional crosswalk is provided in **Table S5**.

Spatial research effort was calculated at the article level. An article represented in one region received weight 1; an article spanning *k* regions received weight 1/*k* in each, so every article contributed a total weight of 1. For region r, observed effort was the regional sum of fractional article weights divided by total fractional effort, and expected effort was the region’s proportion of mapped jaguar-range area. We defined the research concentration index as:

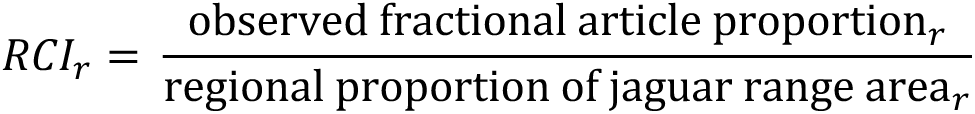

Values >1 indicated greater article-based effort than expected from area and values <1 lower representation; this was an area-based reference rather than a prescription for proportional research allocation.

Uncertainty was quantified with 1,000 bootstrap replicates in which articles were resampled with replacement. Regions were classified as concentrated when the lower 95% confidence limit exceeded 1, underrepresented when the upper limit was below 1, and uncertain when the interval overlapped 1. Spatial-scale sensitivity repeated the analysis for the 15 subregions, and scope sensitivity used the inclusive corpus. Ecoregion-level fractional effort per 10,000 km² was calculated descriptively for ecoregion fragments containing at least 10,000 km² of mapped range. Broad-region, subregion, ecoregion, bootstrap, and sensitivity outputs are reported in **Table S6**. We also compared the temporal and thematic composition of the georeferenced terrestrial subset with the remainder of the primary corpus (**Table S7**).

### 2.5. Anthropogenic context and candidate spatial knowledge gaps

Contemporary anthropogenic context was represented by the Wildlife Conservation Society Human Impact Index (HII) annual layer for 2020 (Venter *et al*., 2016a, 2016b), downloaded on 17 October 2025. Raw raster values were divided by 100 to recover the intended 0–64 scale and processed with terra (Hijmans *et al*., 2022). We evaluated HII at two scales: coverage-weighted summaries across each broad region within the jaguar range and mean HII within 25-, 50-, and 100-km buffers around georeferenced study locations, with 50 km specified as the primary scale. To prevent studies with many coordinates from dominating regional estimates, coordinate-level HII values were first averaged within each article–region combination. We then calculated the difference between mean study-location HII and the corresponding region-wide mean.

Spearman’s rank correlations evaluated associations between regional RCI and (1) region-wide mean HII and (2) the difference between study-location and region-wide HII. These analyses were descriptive and did not treat HII as a direct measure of jaguar population status, conflict intensity, ecological response, or conservation priority. HII product metadata, regional and study-location summaries, association tests, and buffer-radius sensitivity are reported in **Table S8**. Candidate spatial knowledge gaps combined RCI uncertainty with regional HII. Regions represented by fewer than five linked articles were classified as having insufficient evidence. Underrepresentation required the upper 95% bootstrap confidence limit of RCI to be <1. Underrepresented regions with mean HII at or above the median across the nine broad regions were classified as higher-HII candidate gaps; those below the median as candidate gaps. These profiles were intended as transparent screening categories rather than evidence of a universal mismatch between research effort and conservation needs; complete decision rules and regional profiles are provided in **Table S8**. Exploratory regional research-domain coverage was evaluated for broad regions represented by at least ten linked articles (**Table S9**), and the temporal, thematic, and geographic representation of conservation-genetics studies is reported in **Table S10**. All analyses were conducted in R 4.5.1 (R Core Team, 2026); package versions and analytical roles are listed in **Table S4**.

## 3. RESULTS

### 3.1. Analytical corpus and publication growth

The database searches retrieved 2,702 records, of which 1,169 were duplicates. Screening of the remaining 1,533 records excluded 505, resulting in a source dataset of 1,028 articles (**Figure S1**; **Table S1**). The metadata-based scope assessment classified 683 records as high-confidence jaguar focus, 174 as comparative or multispecies jaguar focus, 132 as contextual or uncertain, 36 as indexed-keyword-only or likely incidental, and three as having no direct jaguar signal. The primary analytical corpus therefore comprised 857 articles (83.4% of the source dataset), whereas the inclusive sensitivity corpus comprised 989 (96.2%; **Figure 1A**; **Table S1**).

**Figure 1.**
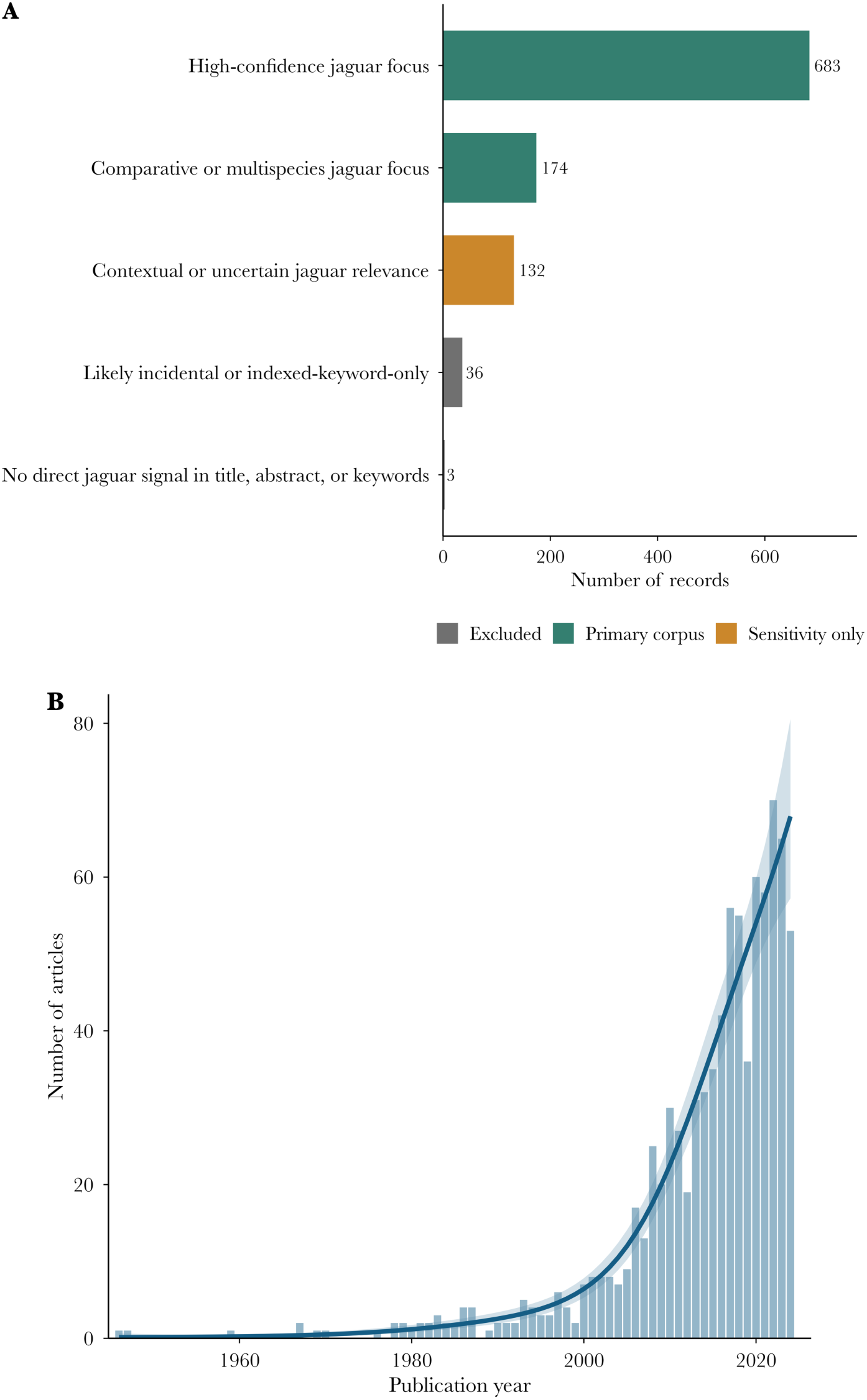
Corpus definition and growth of jaguar research. **(A)** Metadata-based analytical-scope classification used to define the primary corpus, the sensitivity-only records, and excluded records. **(B)** Annual numbers of articles in the primary analytical corpus from 1946 to 2024. The line shows the fitted negative-binomial generalized additive model and the band shows its 95% confidence interval.

Publication output increased markedly from 1946 to 2024 (**Figure 1B**). Annual output remained at nine or fewer articles through 2005, exceeded ten for the first time in 2006, and peaked at 70 articles in 2022. Overall, 92.3% of articles were published from 2000 onward and 78.1% from 2010 onward. The negative-binomial generalized additive model provided the best description of this trajectory (AIC = 285.51), outperforming both the linear (AIC = 292.07) and intercept-only models (AIC = 481.68), and explained 96.0% of deviance (adjusted R² = 0.957; Table S4). Annual publication patterns were nearly identical between the primary and inclusive corpora (Spearman’s ρ = 0.99, n = 79, p < 0.001; **Figure S2**; **Table S4**).

### 3.2. Research domains, methodological approaches, and keyword structure

Research was unevenly distributed among the predefined domains and methodological approaches (**Figure 2**; **Table S2**). Population and abundance were the most frequent research-domain indicator (85 articles; 9.9%), followed by human–wildlife conflict and coexistence (77; 9.0%), movement and connectivity (50; 5.8%), diet and trophic ecology (42; 4.9%), and health, disease, and pathogens (40; 4.7%). The remaining domains each occurred in fewer than 4% of articles. Camera trapping was the most frequent methodological indicator (86 articles; 10.0%), whereas each of the other five methodological categories occurred in 3.2% or less of the primary corpus. Broader supported classifications increased category frequencies but retained the same general ranking (**Table S2**).

**Figure 2.**
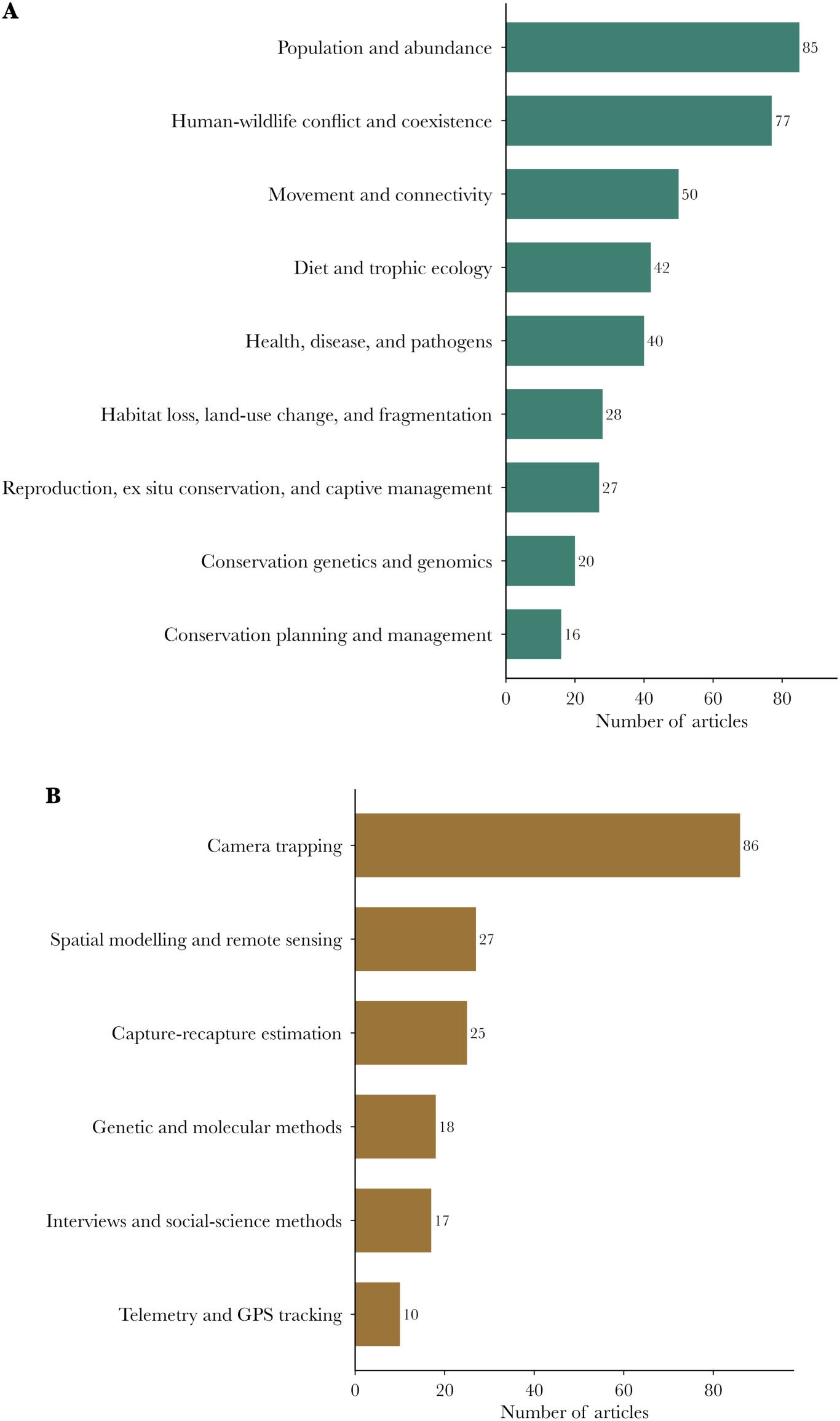
Research domains and methodological approaches. **(A)** Frequencies of high-specificity research-domain indicators. **(B)** Frequencies of high-specificity methodological indicators. Indicators required evidence in the title or author keywords, and an article could contribute to more than one category.

After keyword standardization, the most frequent author keywords were “puma” (98 articles; 11.4%), “camera trapping” (72; 8.4%), “conservation” (53; 6.2%), “human–wildlife conflict” (38; 4.4%), and “predation” (27; 3.2%; **Figure S3**; **Table S3**). The filtered author-keyword network contained 13 nodes and 26 edges and resolved three communities (modularity = 0.363; **Figure S4**; **Table S3**). One connected camera trapping, conservation, corridor, activity patterns, coexistence, and habitat use; a second grouped puma, human– wildlife conflict, and livestock depredation; and a third linked Pantanal, livestock, home range, and predation. “Puma” had the highest degree and Jaccard-weighted strength, whereas “conservation” had the highest betweenness centrality. The sensitivity network combining author and indexed keywords also resolved three communities, although with lower modularity (Q = 0.128; **Table S3**).

### 3.3. Temporal composition of research domains and methodological approaches

Temporal analyses included 824 articles published from 1990 onward. Although several research-domain smooth terms had low unadjusted p-values, none retained strong evidence after Benjamini–Hochberg adjustment (**Figure S5**; **Table S4**). The lowest adjusted values occurred for population and abundance and human–wildlife conflict and coexistence (both *p_BH_* = 0.085); all other domains had *p_BH_* ≥ 0.129. Likewise, none of the fitted methodological indicators showed an adjusted temporal association (all *p_BH_* = 0.322; **Figure S6**; **Table S4**). Telemetry and GPS tracking was not modelled because it did not meet the minimum sample-size criterion.

### 3.4. Spatial coverage and concentration of research

Of 1,762 coordinate rows in the source database, 1,117 were retained after validation and successfully linked to bibliographic records. Within the primary corpus, 673 coordinate rows were assigned to terrestrial broad regions within the mapped jaguar range, representing 237 unique articles (27.7% of the primary corpus; **Table S5**). This georeferenced subset was more strongly represented by articles from the 2010s and by population-and-abundance studies than the remainder of the corpus; differences for other research domains were comparatively small (**Table S7**).

Article-based research effort differed substantially from the area-proportional expectation across the nine broad regions (**Figure 3**; **Table 1**; **Table S6**). The Pantanal showed the highest research concentration index (RCI = 9.89, 95% CI = 6.68–13.20), followed by Mesoamerica (RCI = 8.91, 95% CI = 7.55–10.19) and the Atlantic Forest (RCI = 3.61, 95% CI = 2.44–4.79); all three were classified as concentrated. Cerrado and Caatinga, Chaco and Chiquitano dry forests, and Llanos and Orinoco had confidence intervals overlapping one and were therefore classified as uncertain. In contrast, the Andes and Chocó–Darién (RCI = 0.52, 95% CI = 0.25–0.86), Amazon (RCI = 0.22, 95% CI = 0.15–0.29), and Guiana Shield (RCI = 0.21, 95% CI = 0.07–0.36) were underrepresented. The contrast was particularly pronounced in the Amazon, which contained 51.8% of the mapped jaguar-range area but only 11.2% of fractional article effort (**Figure 3**; **Table 1**).

**Figure 3.**
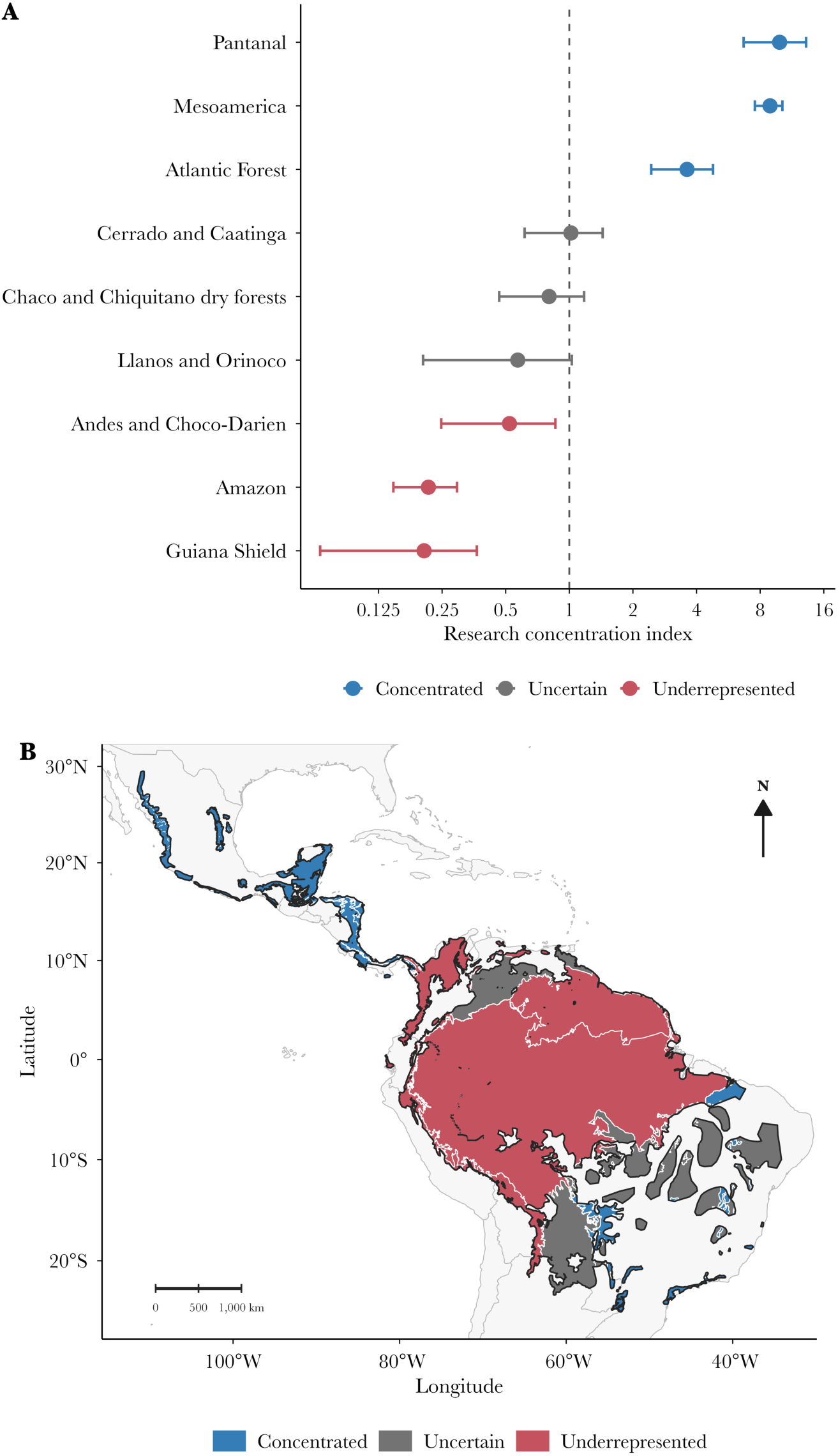
Spatial concentration of jaguar research. **(A)** Research concentration index for each broad region, calculated as observed fractional article effort divided by effort expected from regional area within the mapped jaguar range. Horizontal bars are 95% article-bootstrap confidence intervals. **(B)** Geographic distribution of the corresponding evidence categories. Point estimates above one indicate concentration relative to area, whereas values below one indicate lower estimated representation. Evidence categories were assigned according to whether the 95% bootstrap confidence interval excluded one.

**Table 1.** Broad-region distribution of georeferenced jaguar research relative to mapped range area.

| Broad region | Articles | Range area (%) | Research effort (%) | RCI (95% CI) | Evidence category |
| --- | --- | --- | --- | --- | --- |
| Pantanal | 35 | 1.2 | 11.5 | 9.89 (6.68–13.20) | Concentrated |
| Mesoamerica | 104 | 4.7 | 42.1 | 8.91 (7.55–10.19) | Concentrated |
| Atlantic Forest | 33 | 3.2 | 11.5 | 3.61 (2.44–4.79) | Concentrated |
| Cerrado and Caatinga | 28 | 8.3 | 8.5 | 1.02 (0.61–1.44) | Uncertain |
| Chaco and Chiquitano dry forests | 22 | 8.6 | 6.9 | 0.80 (0.47–1.17) | Uncertain |
| Llanos and Orinoco | 8 | 4.2 | 2.4 | 0.57 (0.20–1.03) | Uncertain |
| Andes and Chocó–Darién | 13 | 7.0 | 3.6 | 0.52 (0.25–0.86) | Underrepresented |
| Amazon | 34 | 51.8 | 11.2 | 0.22 (0.15–0.29) | Underrepresented |
| Guiana Shield | 10 | 11.0 | 2.3 | 0.21 (0.07–0.36) | Underrepresented |

Article counts are unique within regions and may sum to more than 237 because articles spanning multiple regions were counted in each represented region. Research effort was calculated using fractional article weights so that each article contributed a total weight of one across regions. RCI = research concentration index. Evidence categories were based on whether the 95% bootstrap confidence interval excluded 1.

The broad-region pattern was robust to corpus definition: concentration estimates from the primary and inclusive corpora were strongly correlated (Spearman’s ρ = 0.98, n = 9, *p* < 0.001), and all regions retained the same evidence category (**Figure S7**; **Table S6**). Analyses at the 15-subregion scale similarly identified strong concentration in Central American subregions and underrepresentation in the Amazon, Guiana Shield, Andes and inter-Andean valleys, and Chocó–Darién and Pacific forests (**Table S6**). At the ecoregion scale, 19 of 71 ecoregions containing at least 10,000 km² of mapped jaguar range had no linked article (**Figure S8**; **Table S6**).

### 3.5. Anthropogenic context and candidate spatial knowledge gaps

Region-wide mean HII ranged from 1.54 in the Guiana Shield and 1.88 in the Amazon to 7.25 in the Andes and Chocó–Darién and 8.28 in the Atlantic Forest (**Figure 4A**; **Table S8**). Research concentration was not significantly associated with region-wide mean HII (Spearman’s ρ = 0.40, n = 9, *p* = 0.286; **Figure 4A**; **Table S8**) or with the difference between study-location and region-wide HII (Spearman’s ρ = 0.05, n = 9, *p* = 0.898; **Table S8**). Nevertheless, mean HII within 50 km of study locations exceeded the corresponding regional mean in seven of nine regions (**Figure 4B**; **Table S8**). Relative to each region’s HII interquartile range, study locations were concentrated in higher-HII portions of the Atlantic Forest, Amazon, and Guiana Shield, whereas the remaining six regions fell within their regional distributions. These patterns were similar across the 25-, 50-, and 100-km buffer analyses (**Figure S9**). Combining research concentration with regional HII identified the Andes and Chocó–Darién as a higher-HII candidate spatial knowledge gap and the Amazon and Guiana Shield as candidate gaps (**Table 2**; **Table S8**). The Atlantic Forest and Mesoamerica represented research concentrations partly aligned with higher HII, whereas the Pantanal combined high research concentration with comparatively low regional HII. No clear gap signal was identified for Cerrado and Caatinga, Chaco and Chiquitano dry forests, or Llanos and Orinoco because their RCI confidence intervals overlapped one.

**Figure 4.**
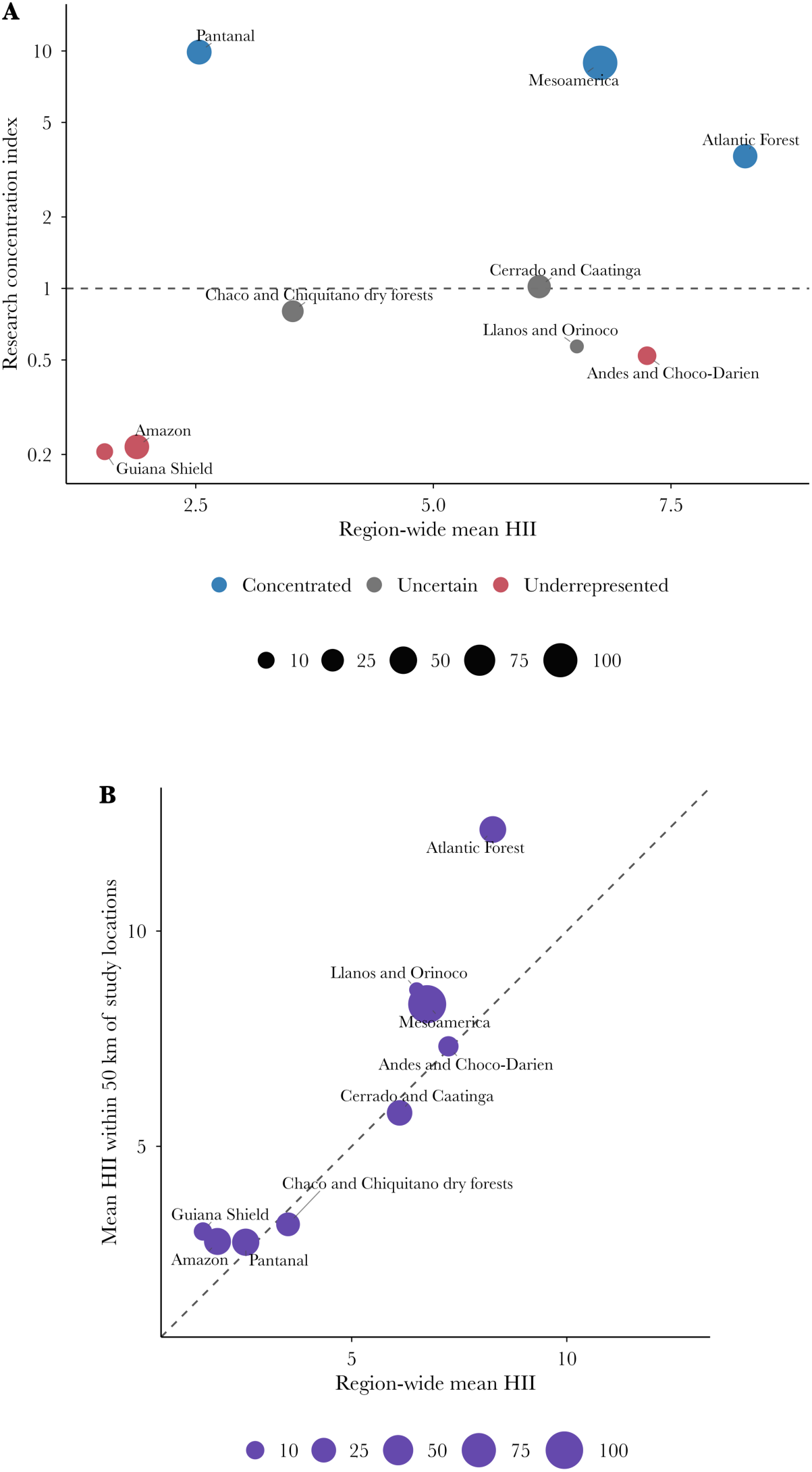
Research concentration and contemporary anthropogenic context. **(A)** Research concentration index in relation to region-wide mean Human Impact Index. **(B)** Mean HII within 50 km of study locations relative to region-wide HII availability. The comparisons represent conditions in 2020.

**Table 2.** Candidate spatial knowledge gaps and anthropogenic context.

| Broad region | Articles | RCI (95% CI) | Regional mean HII | Study-location context | Evidence profile |
| --- | --- | --- | --- | --- | --- |
| Andes and Chocó–Darién | 13 | 0.52 (0.25–0.86) | 7.25 | Within regional IQR | Higher-HII candidate gap |
| Amazon | 34 | 0.22 (0.15–0.29) | 1.88 | Higher-HII portions | Candidate gap |
| Guiana Shield | 10 | 0.21 (0.07–0.36) | 1.54 | Higher-HII portions | Candidate gap |

RCI = research concentration index; HII = Human Impact Index; IQR = interquartile range. Candidate gaps were underrepresented regions whose upper 95% RCI confidence limit was below 1. Higher-HII candidate gaps additionally had regional mean HII at or above the median across the nine broad regions.

### 3.6. Exploratory regional domain coverage and conservation genetics

Exploratory comparisons provided little evidence of strong regional differentiation in research-domain coverage. Only two region–domain combinations exceeded the corresponding proportion in the full georeferenced subset: habitat loss, land-use change, and fragmentation in the Chaco and Chiquitano dry forests, and reproduction, ex situ conservation, and captive management in the Pantanal (**Figure S10**; **Table S9**). No other regional comparison differed clearly from the reference proportion. Conservation genetics and genomics was represented by 20 high-specificity articles (2.3% of the primary corpus), published between 2001 and 2024 (**Figure S11**; **Table S10**). Only eight were represented in the terrestrial spatial subset, with most linked to Mesoamerica and no georeferenced genetics articles in the Andes and Chocó–Darién, Chaco and Chiquitano dry forests, or Llanos and Orinoco (**Figure S12**; **Table S10**).

## 4. DISCUSSION

Our results reveal a rapidly expanding but geographically uneven body of jaguar research. The research-expansion and knowledge-structure hypotheses were supported by strong nonlinear growth in publication output, unequal representation of research domains and methods, and identifiable author-keyword communities. The spatial-concentration hypothesis was also supported, with article-based effort departing markedly from the area-proportional expectation and remaining robust to the broader corpus definition. By contrast, the anthropogenic-alignment hypothesis was not supported at the regional scale. Together, these findings indicate a structured and spatially concentrated evidence base, but not one that can be characterized simply as fragmented or universally mismatched with conservation conditions.

### 4.1. Rapid expansion and the methodological development of jaguar research

The acceleration of jaguar publications after 2000 parallels the broader expansion of carnivore and conservation research. Large-bodied and geographically widespread carnivores tend to attract relatively high scientific attention, although publication effort is not necessarily proportional to extinction risk or conservation need (Brooke *et al*., 2014). Jaguars combine several characteristics associated with scientific visibility, including large body size, broad distribution, cultural prominence, frequent interactions with people, and use as a focal species for landscape conservation. Publication growth therefore indicates maturation of a research field centered on a charismatic and management-relevant carnivore but does not by itself demonstrate that the resulting evidence is geographically representative or sufficient for range-wide conservation.

Camera trapping was the most frequent explicitly identified methodological approach. Its prominence is consistent with its expansion across mammalian research and its suitability for individually recognizable felids (McCallum, 2013). For jaguars, systematic camera trapping enabled standardized abundance and density estimation across multiple environments and became a central component of population monitoring (Silver *et al*., 2004). However, methodological prevalence should not be interpreted as evidence that camera trapping caused the overall increase in publication output, because our analyses did not test that relationship.

Methodological coverage also differs from temporal depth. Many surveys provide valuable snapshots of occurrence, abundance, or density, whereas repeated long-term monitoring remains less common. Camera-trap density estimates can be sensitive to sampling design and analytical assumptions (Tobler & Powell, 2013), while long-term work in Belize showed that detection, residency, sex ratios, and temporary emigration vary through time in ways difficult to detect from isolated surveys (Harmsen *et al*., 2017). Continued development should therefore emphasize repeated and comparable designs, explicit treatment of detection processes, and integration of camera trapping with telemetry, genetics, demographic models, and social information.

### 4.2. A structured, management-oriented, but not strongly fragmented evidence base

Population and abundance, human–wildlife conflict and coexistence, and movement and connectivity were the most frequent research-domain indicators. These topics correspond directly to information needed to assess population status, identify functional corridors, understand space use, and manage persistence in shared landscapes. Their prominence may therefore reflect legitimate scientific and conservation demand rather than an undesirable thematic imbalance. More broadly, research effort across Carnivora reflects biological traits, detectability, range size, interactions with people, and human interest as well as conservation status (Brooke *et al*., 2014).

The strong representation of conflict and coexistence is consistent with broader research on large carnivores. Reviews of *Panthera* and other carnivores have documented substantial attention to livestock depredation, attitudes, spatial relationships between people and predators, compensation, and mitigation, alongside persistent geographic and conceptual biases (Krafte Holland *et al*., 2018; Lozano *et al*., 2019; Venumière-Lefebvre *et al*., 2022). A jaguar-specific scoping review similarly identified a substantial Latin American literature on human–jaguar conflict (Solano *et al*., 2026). However, our temporal models did not provide strong adjusted evidence that conflict, population, or other domains increased proportionally through time. Frequency and temporal change are distinct. A domain can be prominent in the cumulative literature without showing a supported directional increase in its annual representation. The most conservative interpretation is therefore that the field expanded broadly while proportional thematic composition remained comparatively stable or too variable to resolve confidently.

Habitat loss, land-use change, and fragmentation were identified less frequently than population, conflict, or movement. This should not be interpreted as evidence that habitat transformation is unimportant in jaguar research. Our high-specificity indicators required explicit evidence in titles or author keywords and could miss studies in which habitat change was treated as background context, a covariate, or an indirect component of distribution, movement, or population analyses. Similar patterns have been reported across Felidae, where habitat loss and fragmentation are often addressed indirectly and research effort remains uneven among topics and analytical approaches (Zanin *et al*., 2015). The frequencies reported here thus describe the explicit organization of bibliographic metadata rather than the complete substantive content of every article.

The keyword network likewise suggests organization without strong compartmentalization. Three communities connected population-monitoring and conservation terms, conflict-related terms, and a Pantanal–livestock–home-range–predation group, while moderate modularity and the weaker modularity of the sensitivity network indicated substantial links among them. The high connectivity of “puma” likely reflects a long tradition of comparative work on sympatric jaguars and pumas, whereas the high betweenness of “conservation” shows that management-oriented concepts bridge otherwise distinct topics. The evidence base is therefore better described as containing identifiable but interconnected research communities than as being divided into isolated thematic domains.

Some areas, including conservation planning, conservation genetics and genomics, and several methodological approaches, were explicitly represented less often, but these frequencies should not be interpreted as formal priority rankings. The small genetics subset, in particular, prevented strong spatial inference, despite the importance of genetic evidence for connectivity, isolation, demographic history, and fragmentation. A useful direction is therefore greater integration of genetics, movement, population monitoring, health, and human dimensions within coordinated and repeated programmes rather than expansion of each component in isolation.

### 4.3. Research concentration reflects both accumulated capacity and uneven geographic coverage

The strongest spatial result was the concentration of article-based research in the Pantanal, Mesoamerica, and Atlantic Forest. This pattern was robust to the operational definition of the corpus, with nearly identical concentration estimates and unchanged regional classifications under the inclusive sensitivity analysis. These regions contain influential field programmes, established institutions, long monitoring histories, or coordinated conservation initiatives. Foundational camera-trap and telemetry studies in the Pantanal, standardized monitoring in Mesoamerica, and multi-institutional population assessments in the Atlantic Forest illustrate how sustained research capacity can generate long-term demographic and management-relevant evidence (Silver *et al*., 2004; Soisalo & Cavalcanti, 2006; Paviolo *et al*., 2016; Harmsen *et al*., 2017).

High concentration should therefore not be interpreted automatically as excessive or inefficient allocation. Repeated work can generate demographic time series, evaluate interventions, improve methodology, and detect population change in ways that isolated surveys cannot. This distinction is especially important in highly threatened systems. In the Atlantic Forest, for example, intensive research coincides with severe habitat loss, fragmentation, population isolation, and anthropogenic mortality, and has documented jaguar persistence in small, fragmented subpopulations (Paviolo *et al*., 2016).

At the same time, concentration in established research regions leaves extensive portions of the species’ distribution comparatively weakly represented. The Amazon contained more than half of the mapped jaguar-range area but only a small share of area-proportional article effort, while the Guiana Shield and Andes and Chocó–Darién were also underrepresented after bootstrap uncertainty was considered. This pattern is consistent with broader evidence that conservation knowledge tends to accumulate in accessible, institutionally connected, and economically favored locations (Trimble & van Aarde, 2012; Amano & Sutherland, 2013; Wilson *et al*., 2016; Lessa *et al*., 2019).

Such underrepresentation matters because ecological relationships, population densities, threats, prey communities, landscape resistance, and responses to people are unlikely to be constant across the Neotropics. Estimates or management recommendations derived from well-studied floodplains, protected tropical forests, or fragmented Atlantic Forest landscapes may therefore transfer poorly to Amazonian, Guianan, Andean, or Pacific systems without regional validation. This is consistent with the broader tendency for ecological models to lose predictive performance outside the environments, scales, or sampling conditions in which they were developed (Yates *et al*., 2018).

The area-proportional index nevertheless should not be interpreted as implying equal research effort per unit area. Large, remote, low-density, or relatively intact regions may require different sampling intensity and study designs from smaller or highly transformed landscapes. Its value is to provide an explicit baseline against which the magnitude of geographic concentration can be evaluated. Under this interpretation, the Amazon and Guiana Shield emerge as candidate gaps because of their low representation relative to extent, whereas the Andes and Chocó–Darién emerge as a higher-HII candidate gap because underrepresentation coincided with comparatively high contemporary human influence. These profiles should be combined with population importance, ecological distinctiveness, connectivity, feasibility, local capacity, and specific conservation questions before research priorities are set.

### 4.4. Anthropogenic pressure does not independently explain research geography

Research concentration was not significantly associated with region-wide mean HII, indicating that scientific attention was not distributed simply according to contemporary anthropogenic pressure. This is consistent with broader analyses showing that conservation research is shaped by more than threat or conservation need. Across Carnivora, extinction risk was not a strong predictor of publication effort, whereas body size and range size were more consistently related to research attention (Brooke *et al*., 2014); conservation research more generally is also influenced by economic resources, institutions, accessibility, political conditions, and historical investment (Wilson *et al*., 2016).

The absence of a regional relationship, however, does not imply that anthropogenic context is irrelevant to study placement. Mean HII around study locations exceeded the corresponding regional average in seven of nine regions, and locations in the Atlantic Forest, Amazon, and Guiana Shield occurred disproportionately in higher-HII portions of those regions. This scale-dependent contrast is plausible because land-use and socioeconomic pressures vary strongly within broad landscapes (Elias *et al*., 2026). Large regions with low average HII may still contain research concentrated near roads, settlements, deforestation fronts, protected-area boundaries, or other accessible and management-relevant areas. Comparable clustering of conservation research near roads and research centers has been documented in the Caatinga (Lessa *et al*., 2019).

Two non-exclusive mechanisms may contribute to this pattern. Research may be directed toward modified landscapes because they contain immediate management problems, or the same infrastructure and human presence may facilitate access and reduce field costs. Our analyses could not distinguish these conservation-demand and opportunity-driven mechanisms. Moreover, the lack of association between regional concentration and study-site departure from regional HII indicates that local targeting of modified landscapes was not systematically stronger in regions receiving more research overall.

The evidence therefore does not support a universal mismatch between jaguar research and conservation conditions. Alignment was context dependent: the Atlantic Forest and Mesoamerica combined high research concentration with relatively high HII, the Pantanal was strongly concentrated despite lower average HII, and the Andes and Chocó– Darién combined low representation with high HII. Because HII measures cumulative landscape modification rather than jaguar abundance, population trend, prey depletion, mortality, conflict, governance, or conservation effectiveness, it should be interpreted as one dimension of contemporary landscape context rather than as a direct measure of where research is most needed (Venter *et al*., 2016a).

### 4.5. From identifying patterns to strengthening the evidence base

The practical implication is not to reduce research in concentrated regions. Long-term programmes in the Pantanal, Mesoamerica, and Atlantic Forest provide demographic time series, intervention evaluation, population histories, and methodological consistency that isolated surveys cannot replace. A more effective strategy is additive. Maintain these programmes while expanding representative and locally led research in the Amazon, Guiana Shield, Andes and Chocó–Darién, and other poorly sampled subregions. Recent work in the Brazilian Cerrado illustrates how expert-validated collaborative monitoring can expand spatial coverage into agricultural and private landscapes underrepresented by traditional datasets, providing one practical approach for reducing geographic knowledge gaps (Silva *et al*., 2026).

Expansion should also improve the type and comparability of evidence produced. For conflict and coexistence, this means moving beyond repeated documentation of depredation, attitudes, or correlates toward testing the durability, transferability, costs, and social consequences of mitigation, because interventions often have limited or context-dependent evidence (Krafte Holland *et al*., 2018; Artelle *et al*., 2024). More broadly, shared protocols for camera trapping, genetics, telemetry, health screening, human-dimensions research, and reporting of sampling coordinates would facilitate synthesis without requiring all regions to address identical questions. Range-wide jaguar conservation already depends on linking core populations through transboundary corridors and coordinated planning (Rabinowitz & Zeller, 2010); a similarly connected evidence base would improve the ability to distinguish general relationships from region-specific patterns and to evaluate whether strategies developed in well-studied landscapes transfer across the species’ range.

### 4.6. Limitations and conclusions

Several limitations qualify our conclusions. Web of Science and Scopus do not fully represent regional journals, non-indexed literature, technical reports, theses, government documents, or other grey literature, and unequal database coverage may underrepresent knowledge produced in parts of Latin America and the Caribbean despite the absence of language restrictions (Mongeon & Paul-Hus, 2016; Martín-Martín *et al*., 2018). In addition, the scope filter and research-domain and methodological indicators were based on bibliographic metadata rather than full-text assessment. High-specificity rules reduced false positives but may have omitted studies that addressed a topic without emphasizing it in titles or author keywords, and keyword availability and usage vary among journals, authors, languages, and periods (Aria & Cuccurullo, 2017; Donthu *et al*., 2021).

The spatial analysis was restricted to 237 georeferenced terrestrial articles, and this subset was more recent and more strongly represented by population-and-abundance studies than the remainder of the corpus. Article-based weighting and sensitivity analyses reduced the influence of multi-location studies and corpus definition, but missing coordinates may still have affected regional estimates. Broad spatial units and an area-based expectation also simplify a heterogeneous species range, while the 2020 HII represents contemporary cumulative human influence rather than historical conditions, population status, habitat quality, ecological uniqueness, or conservation urgency. Candidate-gap profiles should therefore be interpreted as screening tools rather than definitive rankings.

Despite these constraints, jaguar research has expanded rapidly and developed an interconnected, management-oriented evidence base with substantial long-term knowledge in several regions. Yet georeferenced evidence remains strongly concentrated relative to the species’ mapped distribution, with clear underrepresentation in the Amazon, Guiana Shield, and Andes and Chocó–Darién. Strengthening range-wide conservation evidence will require maintaining intensive long-term programmes where they provide demographic and management value while expanding durable regional capacity and comparable research in poorly represented landscapes. This combined strategy can improve the geographic representativeness and transferability of jaguar science without sacrificing the temporal continuity and institutional investment required for effective conservation.

## Supporting information

https://zenodo.org/records/21934976

