## Supplementary material for "The knowledge landscape of jaguar science: rapid growth, thematic structure, and regional inequalities": https://zenodo.org/records/21934976

### TEXT NORMALIZATION AND METADATA-BASED SCOPE FILTERING

Bibliographic fields were imported as character values, and empty strings and the values NA, N/A, and NULL were interpreted as missing. Field names were standardized to lowercase snake case, publication years were parsed as numeric values, and unique internal article identifiers were assigned according to source-record order. Titles, abstracts, author keywords, and indexed keywords were converted to lowercase, Unicode accents were transliterated to ASCII where possible, punctuation was replaced by spaces, and repeated whitespace was collapsed. The scope filter searched these fields for multilingual jaguar terms, including jaguar, jaguars, *Panthera onca*, *P. onca*, onca, onca-pintada, onza, onza pintada, yaguar, yaguares, yaguarete, yaguaretes, and tigre americano. Word boundaries were imposed to reduce partial matches, and mentions were counted separately within each metadata field.

Records were classified as high-confidence jaguar focus when at least one jaguar term occurred in the title or author keywords. Records without such evidence but containing at least two abstract mentions were classified as comparative or multispecies jaguar focus. Records containing exactly one abstract mention were classified as contextual or uncertain jaguar relevance. Records containing a jaguar term only in indexed keywords were classified as likely incidental or indexed-keyword-only, and records without a detected signal were classified separately. The main analytical corpus combined the high-confidence and comparative or multispecies categories. The inclusive sensitivity corpus additionally retained contextual or uncertain records. Indexed-keyword-only and no-signal records were excluded from both corpora.

### RESEARCH-DOMAIN AND METHODOLOGICAL DICTIONARIES

Research domains and methodological approaches were represented by predefined, non-exclusive, high-specificity dictionaries. Primary classification required a dictionary match in the title or author-keyword field. A broader supported classification used for auditing and sensitivity purposes was defined as a primary match or concordant evidence in both the abstract and indexed-keyword fields. Evidence restricted to only one of these secondary fields was insufficient for the primary classification. The research-domain dictionary included population and abundance; movement and connectivity; human-wildlife conflict and coexistence; habitat loss, land-use change, and fragmentation; conservation planning and management; conservation genetics and genomics; health, disease, and pathogens; reproduction, ex situ conservation, and captive management; and diet and trophic ecology. The methodological dictionary included camera trapping; telemetry and GPS tracking; genetic and molecular methods; spatial modelling and remote sensing; interviews and social-science methods; and capture-recapture estimation.

Because categories were non-exclusive, an article could be assigned to multiple domains or methods or to none of the predefined categories. A legacy conflict-classification variable retained from the source dataset was preserved for auditing but did not override the standardized rules. The conservation-genetics indicator required specific evidence related to population or landscape genetics, population genomics, genetic diversity, gene flow, genetic connectivity, inbreeding, microsatellites, or single-nucleotide polymorphisms. Generic taxonomic, phylogenetic, pathogen-genomics, or molecular mentions were excluded unless accompanied by an explicit population, landscape, diversity, connectivity, fragmentation, or conservation context.

### AUTHOR-KEYWORD STANDARDIZATION AND NETWORK ANALYSIS

Author-keyword strings were separated at semicolons, converted to lowercase, transliterated where necessary, and standardized for punctuation, hyphenation, and whitespace. Taxonomic search anchors were removed because their frequency resulted directly from the search strategy. Generic bibliographic expressions, broad taxonomic descriptors, and country or continental names were removed using a predefined stop list. Synonymous expressions were harmonized using rule-based transformations and a reviewed synonym table. For example, camera trap variants were standardized as camera trapping, whereas *Puma concolor*, cougar, and mountain lion were standardized as puma. Duplicate occurrences of a standardized keyword within an article were counted once.

The 25 most frequent author keywords were retained for descriptive presentation. Keywords occurring in at least five articles were eligible for network construction, after which the 20 most frequent eligible terms were retained. Every unique unordered keyword pair within an article contributed at most once to the edge count. Edges occurring in fewer than three articles were excluded. The network was represented as an undirected weighted graph using igraph package (v.2.3.3.) For keywords i and j, Jaccard similarity was calculated as:

$$J_{ij} = n_{ij} / (n_i + n_j - n_{ij})$$

where $n_{ij}$ is the number of articles containing both keywords and $n_i$ and $n_j$ are their respective article frequencies. Raw co-occurrence counts and association strength were retained as secondary edge metrics.

The graph two-core was retained when it contained at least six nodes; otherwise, the largest connected component was used. Communities were detected with the Louvain algorithm using Jaccard similarities as edge weights. Node importance was summarized by degree, Jaccard-weighted strength, and normalized betweenness centrality, with inverse Jaccard similarity used as weighted-path distance. The force-directed layout was initialized with the fixed project seed. A sensitivity analysis repeated the procedure after combining author and database-indexed keywords.

### TEMPORAL MODEL SPECIFICATIONS AND DIAGNOSTICS

Annual publication counts were completed for every year between 1946 and 2024, assigning zero to years without records. Three count models were fitted to the same response: an intercept-only negative-binomial model, a negative-binomial model containing centred year as a linear predictor, and a negative-binomial generalized additive model containing a smooth function of year. Negative-binomial generalized linear models were fitted with MASS package (v.7.3–65), and generalized additive models were fitted with mgcv package (v.1.9–4) using restricted maximum likelihood.

The spline-basis dimension for the overall publication GAM was limited according to the number of represented years to avoid excessive flexibility. Candidate models were ranked by Akaike Information Criterion. Annual predictions and 95% confidence intervals were calculated on the link scale and transformed to the response scale. Evaluation metrics included effective degrees of freedom, deviance explained, adjusted $R^2$, and lag-one autocorrelation of deviance residuals.

For temporal composition analyses, each article-level primary indicator was fitted from 1990 onward as a binary response with a logit-link GAM: $\text{logit}[P(Y_i = 1)] = \beta_0 + s(\text{year}_i)$. Models were fitted only when an indicator had at least 15 positive observations, at least 15 negative observations, and records spanning at least ten distinct publication years. The maximum basis dimension was seven. Observed proportions were summarized in five-year periods only for visualization; models used article-level binary data. For each domain and methodological model, we recorded effective degrees of freedom, smooth-term probability, deviance explained, AIC, and lag-one autocorrelation of residual annual proportions. Probabilities were adjusted separately within the research-domain and methodological families using the Benjamini-Hochberg false-discovery-rate procedure. The resulting temporal panels were considered exploratory.

### COORDINATE VALIDATION, ARTICLE LINKAGE, AND REGIONALIZATION

The coordinate database contained 1,762 rows. Candidate longitude fields included lon, longitude, long, x, and longitud; candidate latitude fields included lat, latitude, y, and latitud. Numerical validity required longitude between -180 and 180 degrees and latitude between -90 and 90 degrees. When a manual validation field was available, accepted affirmative values were yes, sim, true, 1, valid, and ok. Rows failing numerical or manual validation were excluded. This process retained 1,117 coordinate rows. Exact longitude-latitude duplicates were flagged but not automatically removed because identical coordinates could legitimately refer to distinct articles or sampling events. Retained rows were linked to bibliography records sequentially using a unique source-record identifier, a unique normalized DOI, or a unique normalized article title. A field was used only when it identified one bibliographic record unambiguously. All retained rows were linked successfully.

Coordinates were represented as WGS 84 points. Jaguar distribution was represented by the IUCN Red List digital distribution map associated with Quigley et al. (2017; errata published in 2018). The accompanying IUCN spatial metadata were version 6.3, updated in December 2022, and described the source data as unprojected WGS 84 polygons. Source geometries were validated and dissolved into one external distribution geometry. Area values were not taken from the source SHAPE_Area field because it was expressed in geographic degrees.

Terrestrial ecoregions followed Olson et al. (2001) and were intersected with the jaguar range before area calculation. The generic ecoregion class Lake was treated as non-terrestrial and excluded. A reviewed crosswalk assigned ecoregions to 15 subregions and subsequently to nine broad regions: Mesoamerica; Amazon; Guiana Shield; Andes and Choco-Darien; Llanos and Orinoco; Atlantic Forest; Cerrado and Caatinga; Pantanal; and Chaco and Chiquitano dry forests. Unresolved crosswalk assignments caused the workflow to stop rather than allocate regions automatically. Areas were recalculated after transformation to the World Cylindrical Equal Area projection (EPSG:6933). Final maps were displayed using a locally centred Lambert azimuthal equal-area projection with latitude of origin -10 degrees and central longitude -75 degrees. The final main-corpus spatial subset comprised 673 terrestrial coordinate rows representing 237 unique articles.

### FRACTIONAL ARTICLE EFFORT, CONCENTRATION INDEX, AND BOOTSTRAP UNCERTAINTY

Spatial effort was calculated from unique article-region associations rather than raw coordinate rows. For article $i$, the contribution to each represented region $r$ was:

$$w_{ir} = 1 / k_i$$

where $k_i$ is the number of regions represented by the article. Therefore, the sum of an article's weights across all regions was one, preventing multisite studies and articles with many reported coordinates from receiving disproportionate influence.

For each region, observed effort was the sum of fractional article weights divided by total fractional effort. Expected effort was the proportion of the mapped jaguar-range area contained within the region. The research concentration index was:

$$RCI_r = \text{observed fractional article proportion}_r / \text{regional proportion of jaguar-range area}_r$$

Values above one indicated more article-based effort than expected from area, and values below one indicated comparatively lower representation. This was an explicit reference expectation and not a prescription that research should be allocated strictly by area. Uncertainty was estimated with 1,000 non-parametric bootstrap replicates. Eligible articles were sampled with replacement, and an article selected multiple times contributed according to its bootstrap multiplicity. Fractional effort and concentration indices were recalculated in every replicate. Percentile-based 95% confidence intervals were defined by the 2.5th and 97.5th percentiles.

Regions were classified as concentrated when the lower confidence limit exceeded one, underrepresented when the upper limit was below one, and uncertain or approximately proportional when the interval overlapped one. The complete analysis was repeated for the 15 reviewed subregions and for the inclusive sensitivity corpus. Ecoregion effort per 10,000 km$^2$ was calculated descriptively, with graphical ranking restricted to ecoregion fragments containing at least 10,000 km$^2$ within the jaguar range.

### HUMAN IMPACT INDEX PROCESSING AND EXTRACTION

The raster asset was downloaded from the Wildlife Conservation Society Human Footprint data-access portal on 17 October 2025. The layer represented 2020 conditions in WGS 84 geographic coordinates at approximately 0.002694946-degree resolution. Raw raster values ranged from 0 to 6,400 and were divided by 100 to recover the intended 0–64 scale. The workflow stopped if the rescaled values fell outside this range. Region-wide Human Impact Index (HII) was summarized across the complete portion of each broad region within the jaguar range using coverage-weighted raster extraction with terra (v.1.9–34) and exactextractr packages (v.0.10.1.) Calculated statistics included mean, standard deviation, first quartile, median, third quartile, 90th percentile, minimum, maximum, and the number of intersected cells.

Study-location context was evaluated with circular buffers of 25, 50, and 100 km generated in EPSG:6933 and transformed to the raster coordinate reference system. The 50-km radius was specified as the primary scale. Raster cells were weighted by the proportion of each cell covered by the buffer. Coordinate-level buffer means were averaged within each article-region combination before regional summaries were calculated, preventing articles with many coordinates from dominating the estimates. For each region, we calculated the difference between mean HII around study locations and mean HII available across the region. Study contexts were also classified as below the regional first quartile, within the regional interquartile range, or above the regional third quartile. Spearman rank correlations evaluated research concentration against region-wide mean HII and against the study-location departure from regional mean HII. These associations were treated as descriptive, not causal.

### SCOPE SENSITIVITY AND REPRESENTATIVENESS OF THE SPATIAL SUBSET

Temporal sensitivity compared annual article counts from the 857-record main corpus and the 989-record inclusive corpus using Spearman rank correlation. Spatial sensitivity repeated the broad-region concentration analysis for the inclusive corpus and compared concentration indices, bootstrap intervals, and evidence categories with the main-corpus results. Because only 237 main-corpus articles were represented in the terrestrial spatial subset, we evaluated whether this subset differed from the remaining corpus in temporal and thematic composition. Comparisons included publication-decade proportions and frequencies of the nine primary research-domain indicators. This audit was descriptive; absence of eligible coordinates was not interpreted as absence of research.

### CANDIDATE-GAP SCREENING AND EXPLORATORY REGIONAL DOMAIN COVERAGE

Candidate-gap profiles combined broad-region concentration estimates, bootstrap uncertainty, sample size, and region-wide mean HII. Regions represented by fewer than five linked articles were classified as having insufficient evidence. Underrepresentation required the upper 95% confidence limit of the concentration index to remain below one. Mean HII at or above the median across the nine regions was classified as comparatively high. The decision sequence was: (1) fewer than five linked articles, insufficient evidence; (2) underrepresented with comparatively high HII, higher-priority candidate spatial knowledge gap; (3) underrepresented with HII below the median, candidate spatial knowledge gap; (4) concentrated with comparatively high HII, research concentration partly aligned with higher HII; (5) concentrated with HII below the median, research concentration in a lower-HII region; and (6) all remaining cases, no clear spatial gap signal. These labels were screening categories and not direct rankings of conservation importance. Exploratory regional domain coverage was evaluated only for broad regions containing at least ten linked articles. For each domain, the regional article proportion and Wilson 95% confidence interval were compared with the corresponding proportion in the full georeferenced terrestrial reference set. Relative coverage was expressed as log2[(p_region + 0.01)/(p_reference + 0.01)]. This analysis did not influence candidate-gap classification.

### EXPLORATORY CONSERVATION-GENETICS MODULE

Initial inclusion in the conservation-genetics corpus required the high-specificity primary domain indicator described in section *Research-domain and methodological dictionaries*. Included articles were subsequently classified non-exclusively as population and landscape genetics; genomics and SNP-based studies; mitochondrial and phylogeographic studies; forensic or molecular identification; and hybridization or introgression. Subtype classification used combined normalized title, abstract, author-keyword, and indexed-keyword text. We summarized subtype frequencies, temporal distribution, frequent author keywords, and the proportion of linked articles classified as conservation genetics within each broad region. Because only eight of the 20 metadata-identified genetics articles were represented in the terrestrial spatial subset, all geographic outputs from this module were treated as exploratory.

### COMPUTATIONAL ENVIRONMENT AND REPRODUCIBILITY

All analyses were conducted in R (v.4.5.1). The principal analytical packages and versions are listed in **Table S4**. The final workflow generated record-level scope and indicator audits, coordinate-validation and article-linkage diagnostics, the reviewed regional crosswalk, model outputs, sensitivity analyses, figures, tables, and session information. The complete sessionInfo() output is archived with the analytical materials. Redistribution of source bibliographic records and IUCN spatial data should follow the licensing and use conditions of the original providers.

### SUPPLEMENTARY TABLES

**Table S1.** Search screening and scope.

Available at: https://zenodo.org/records/21934976

**Table S2.** Classification rules and frequencies.

Available at: https://zenodo.org/records/21934976

**Table S3.** Keyword frequencies and networks.

Available at: https://zenodo.org/records/21934976

**Table S4.** Temporal models and software.

Available at: https://zenodo.org/records/21934976

**Table S5.** Spatial audit and regionalization.

Available at: https://zenodo.org/records/21934976

**Table S6.** Research concentration and sensitivity.

Available at: https://zenodo.org/records/21934976

**Table S7.** Spatial subset representativeness.

Available at: https://zenodo.org/records/21934976

**Table S8.** HII context and candidate gaps.

Available at: https://zenodo.org/records/21934976

**Table S9.** Regional domain coverage.

Available at: https://zenodo.org/records/21934976

**Table S10.** Conservation genetics.

Available at: https://zenodo.org/records/21934976

### SUPPLEMENTARY FIGURES

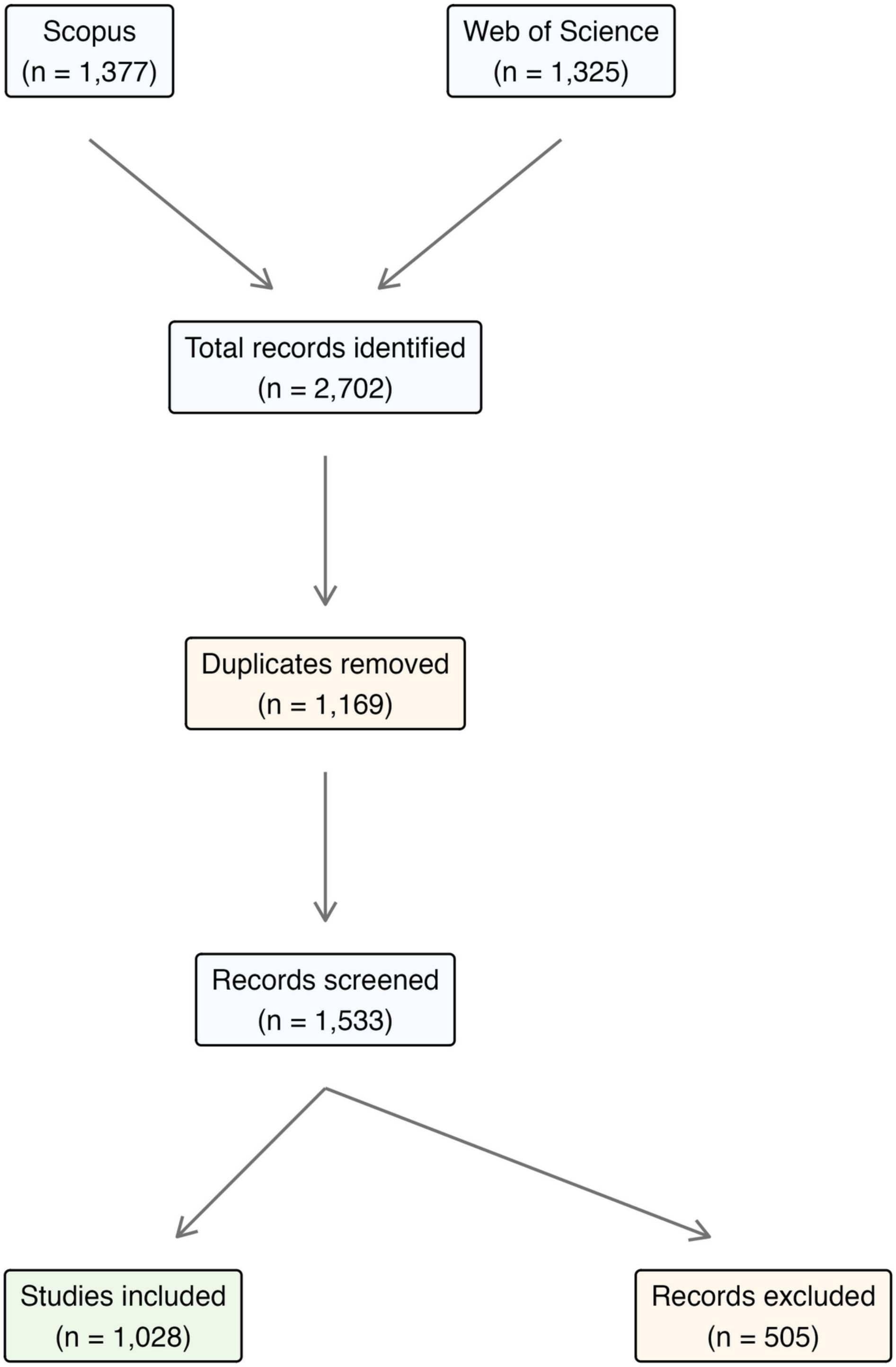


**Figure S1.** PRISMA-style workflow for the bibliographic source dataset. Records retrieved from Scopus and Web of Science were merged, deduplicated, screened, and classified as included or excluded before the metadata-based analytical-scope filter was applied.

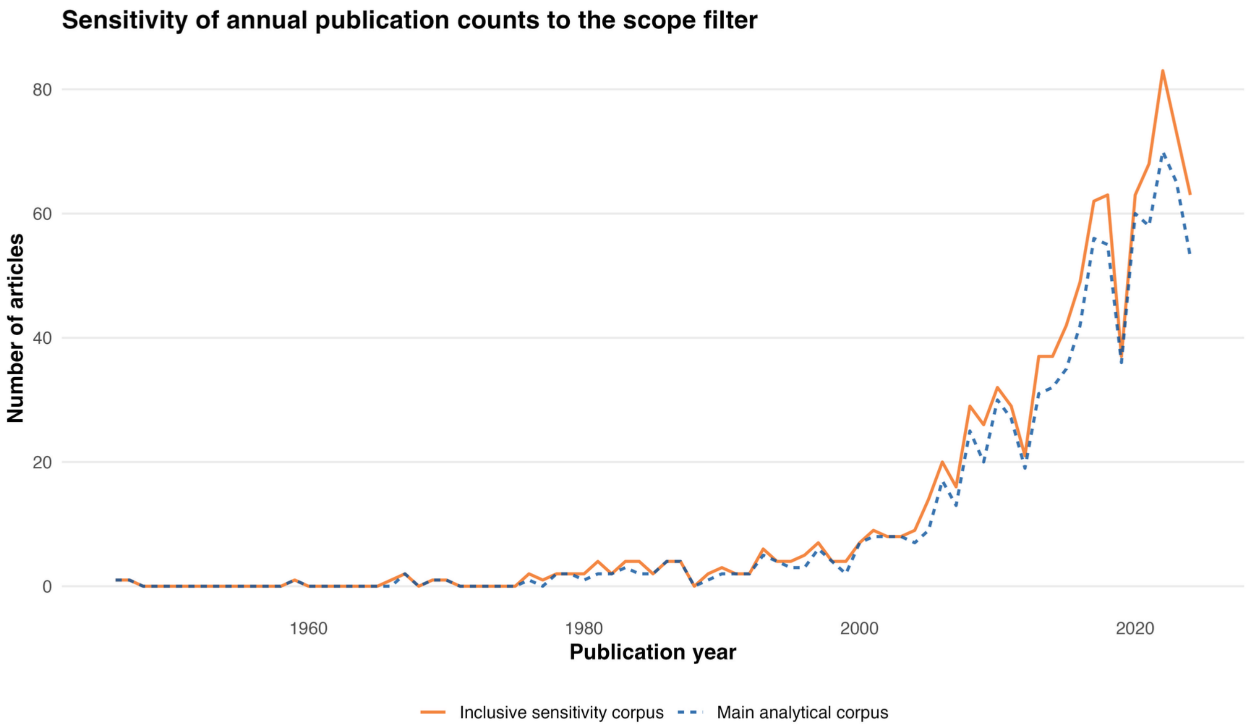


**Figure S2.** Temporal sensitivity to the metadata-based scope definition. Annual publication counts are compared between the primary analytical corpus (n = 857) and the inclusive sensitivity corpus (n = 989). The two annual series were strongly concordant (Spearman's $\rho$ = 0.99).

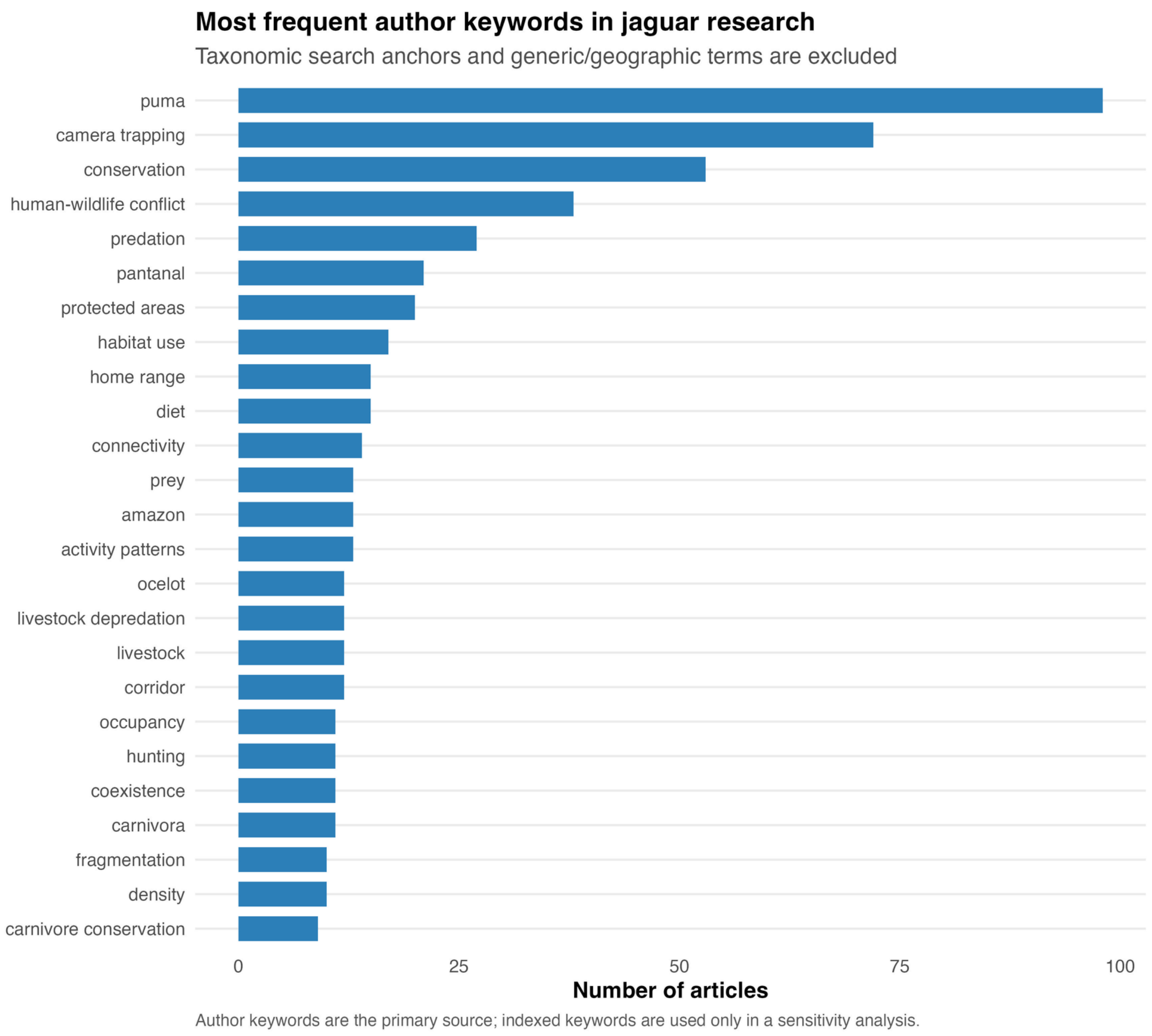


**Figure S3.** Most frequent standardized author keywords in the primary analytical corpus. Taxonomic search anchors, generic bibliographic expressions, broad taxonomic descriptors, and country and administrative geographic terms were excluded, whereas ecologically meaningful regional terms were retained. Each keyword was counted at most once per article.

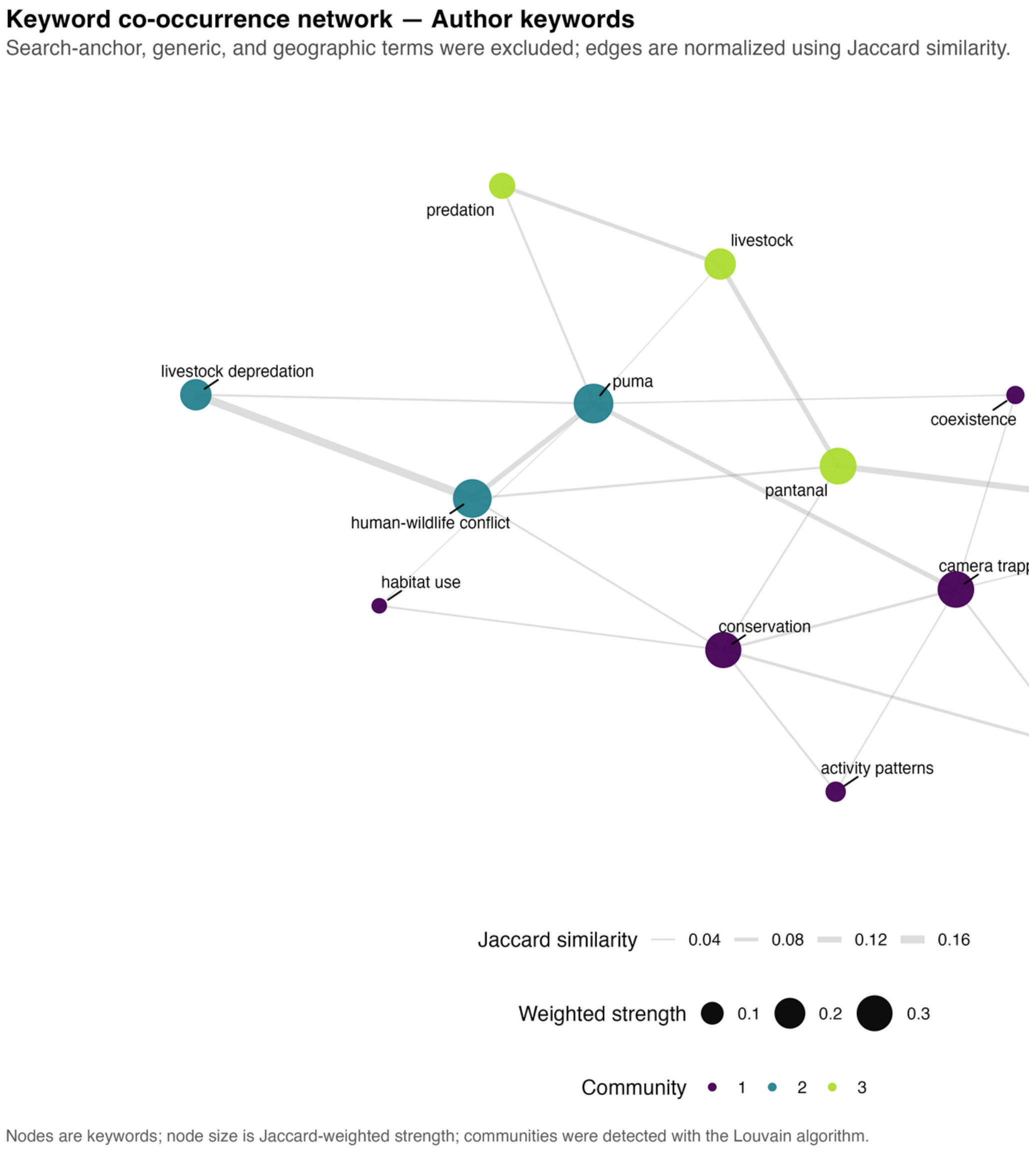


**Figure S4.** Author-keyword co-occurrence network. Nodes represent standardized author keywords, node size represents Jaccard-weighted strength, and node color represents Louvain community membership. Edges represent article-level co-occurrence and are weighted by Jaccard similarity. Search anchors, generic bibliographic expressions, broad taxonomic descriptors, and country and administrative geographic terms were excluded, whereas ecologically meaningful regional terms were retained. The graph two-core was retained when sufficiently large to reduce the influence of peripheral nodes.

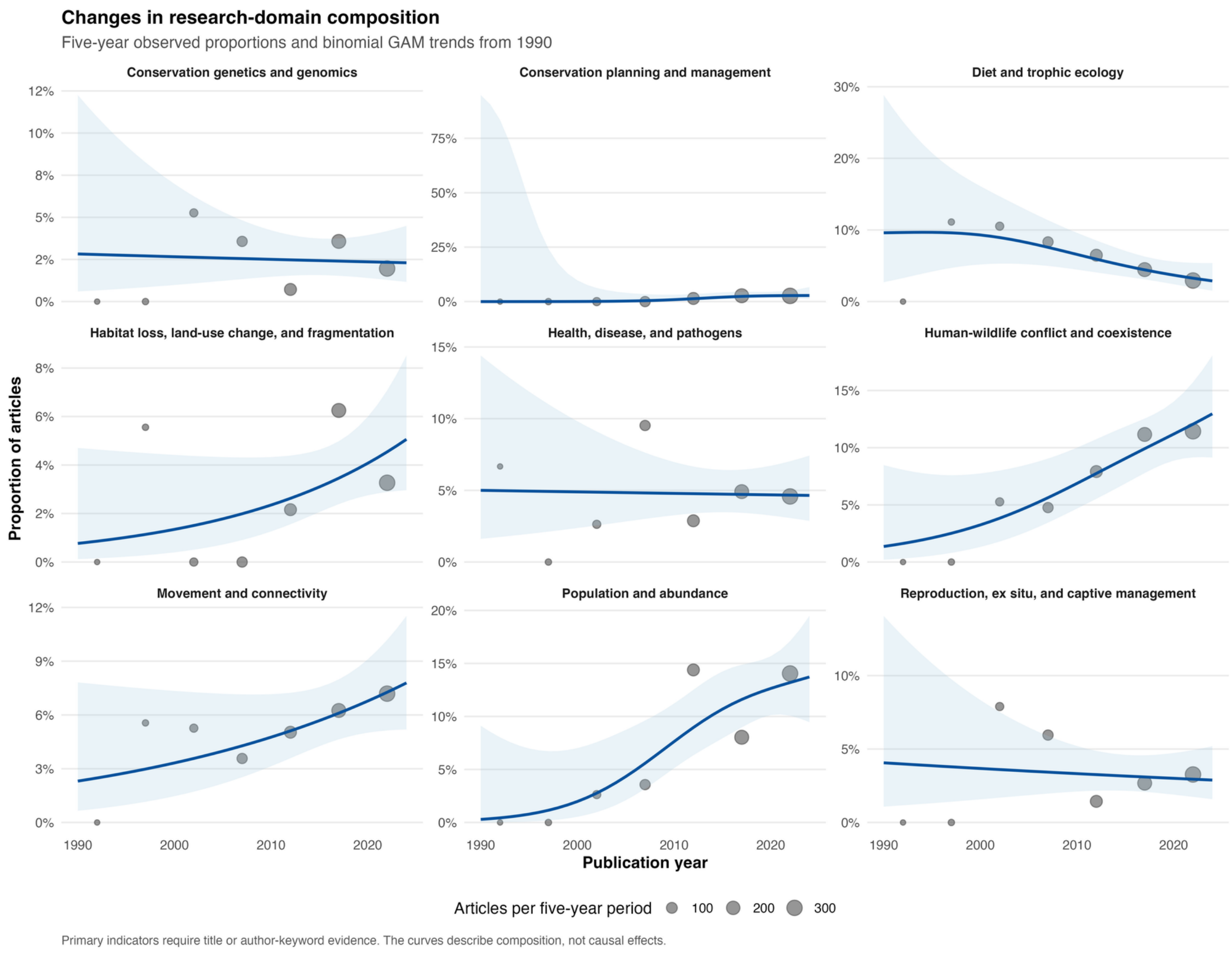


**Figure S5.** Temporal composition of metadata-derived research domains. Points show observed proportions within five-year periods from 1990 onward, with point size proportional to the number of articles in each period. Lines and shaded bands show fitted probabilities and 95% confidence intervals from article-level binomial generalized additive models. Primary indicators required evidence in the title or author-keyword fields. Trends were interpreted as exploratory because none retained strong evidence after Benjamini–Hochberg adjustment.

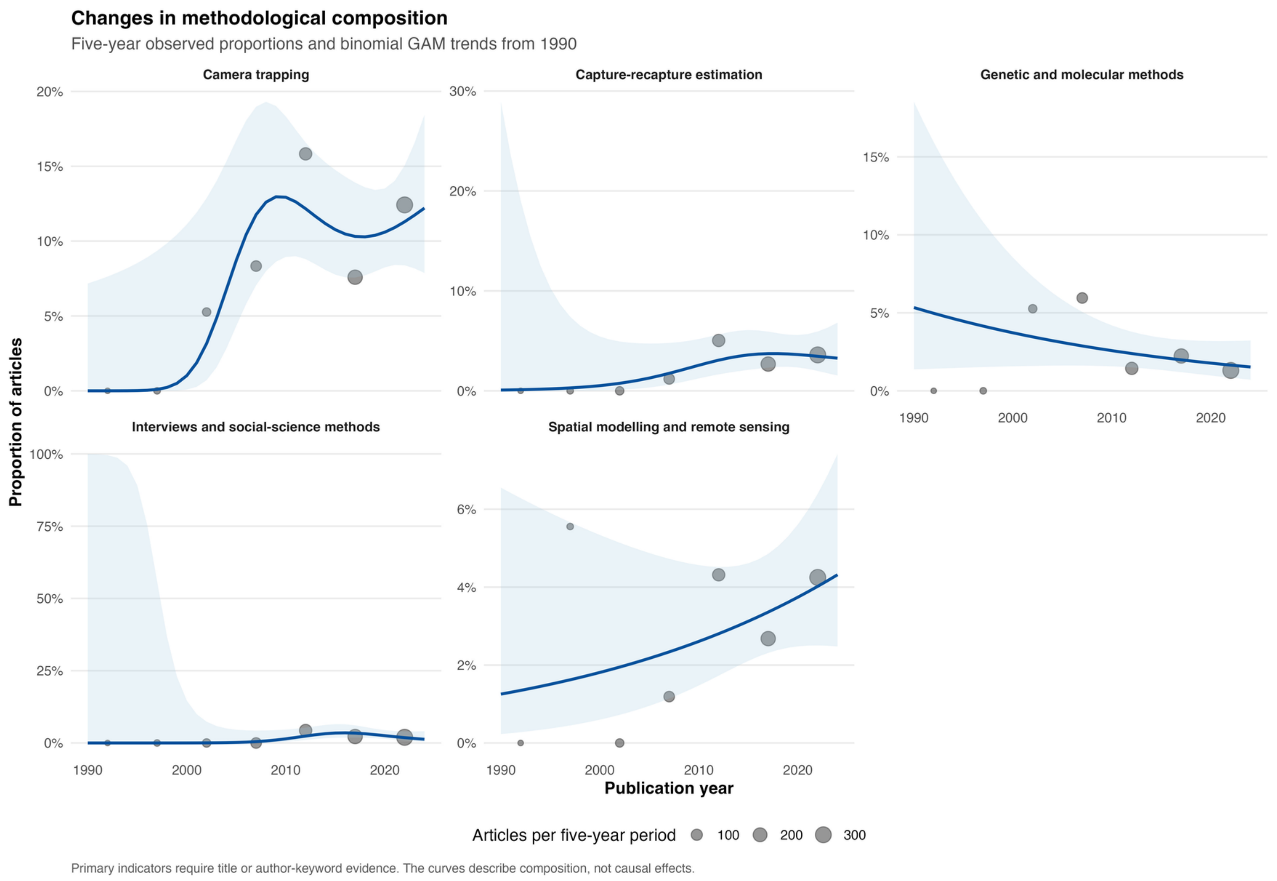


**Figure S6.** Temporal composition of metadata-derived methodological approaches. Points show observed proportions within five-year periods from 1990 onward, with point size proportional to the number of articles in each period. Lines and shaded bands show fitted probabilities and 95% confidence intervals from article-level binomial generalized additive models. Primary indicators required evidence in the title or author-keyword fields. Trends were interpreted as exploratory after Benjamini–Hochberg adjustment.

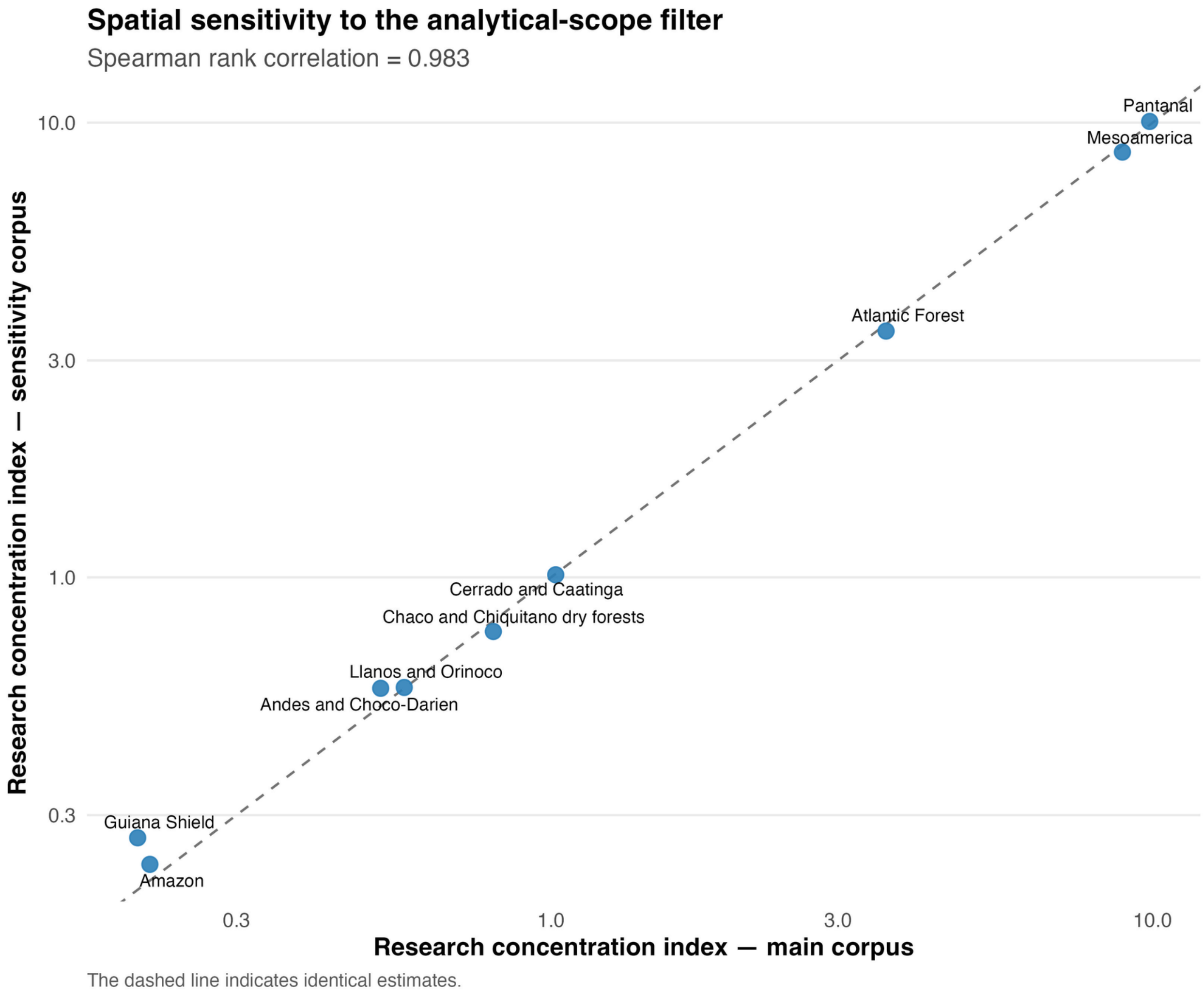


**Figure S7.** Spatial sensitivity to the metadata-based scope definition. Broad-region research concentration indices estimated from the main analytical corpus are compared with indices estimated from the inclusive sensitivity corpus. The dashed line is the 1:1 relationship. Regional rankings were strongly concordant (Spearman rho = 0.983).

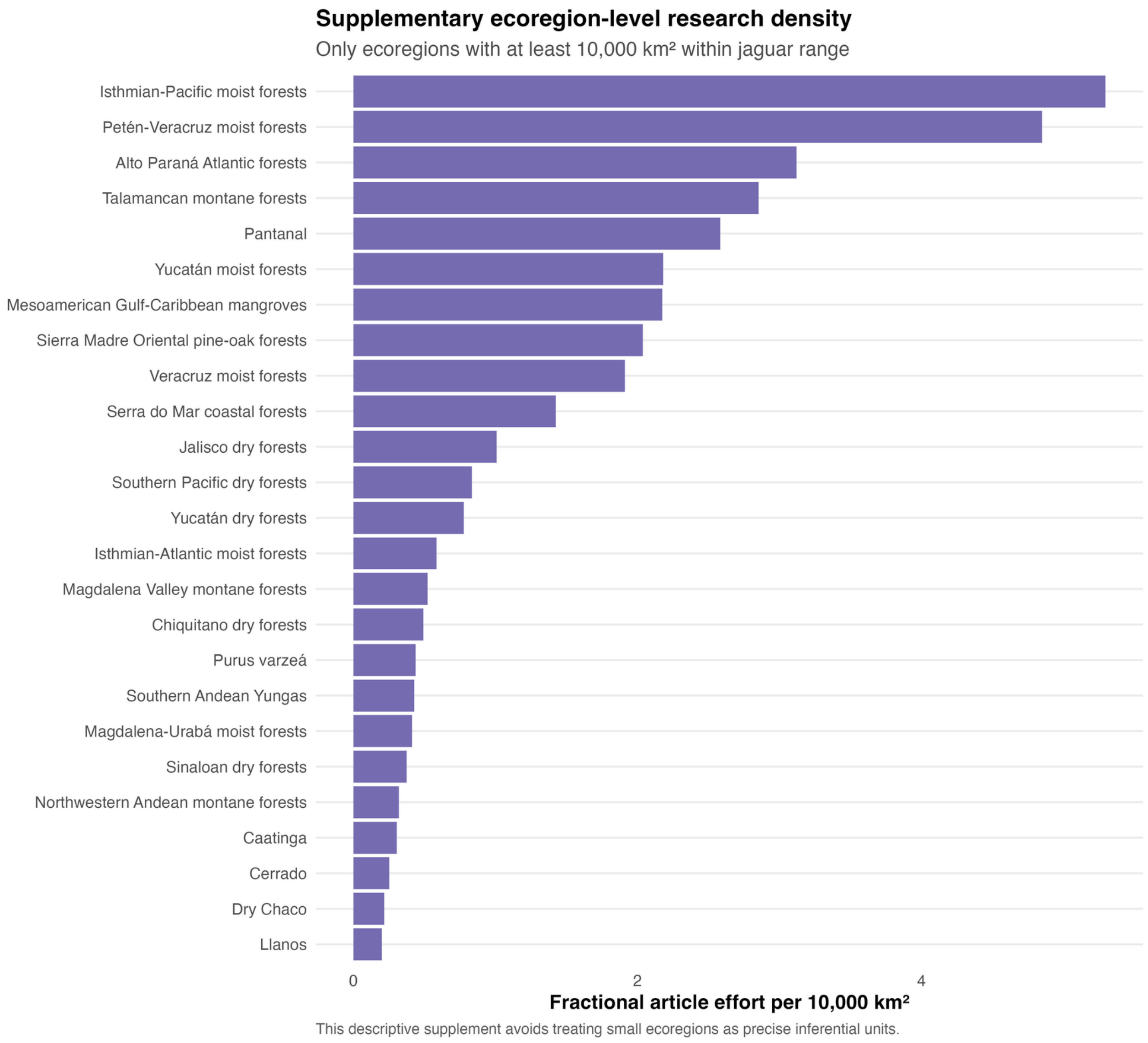


**Figure S8.** Descriptive ecoregion-level research effort. Bars show fractional article effort per 10,000 km2 within the mapped jaguar range. The ranking is restricted to ecoregions containing at least 10,000 km2 within the range to reduce unstable density estimates from small spatial fragments. This figure is descriptive and was not used for primary regional inference.

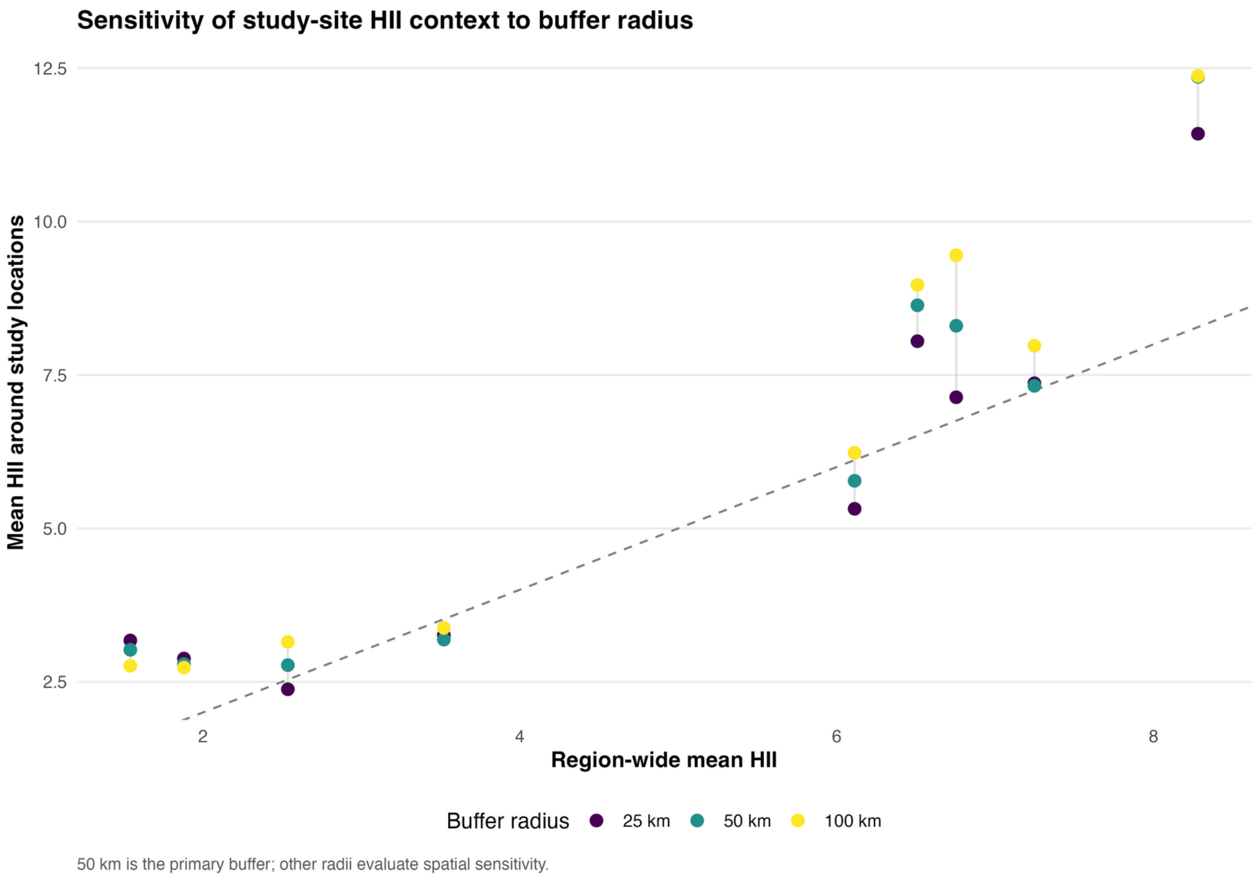


**Figure S9.** Sensitivity of study-location Human Impact Index (HII) to buffer radius. Regional means derived from 25-, 50-, and 100-km buffers around eligible study locations are plotted against region-wide mean HII. The 50-km radius was specified as the primary study-location scale; the other radii were used for sensitivity assessment. The dashed line indicates equality between study-location and region-wide means.

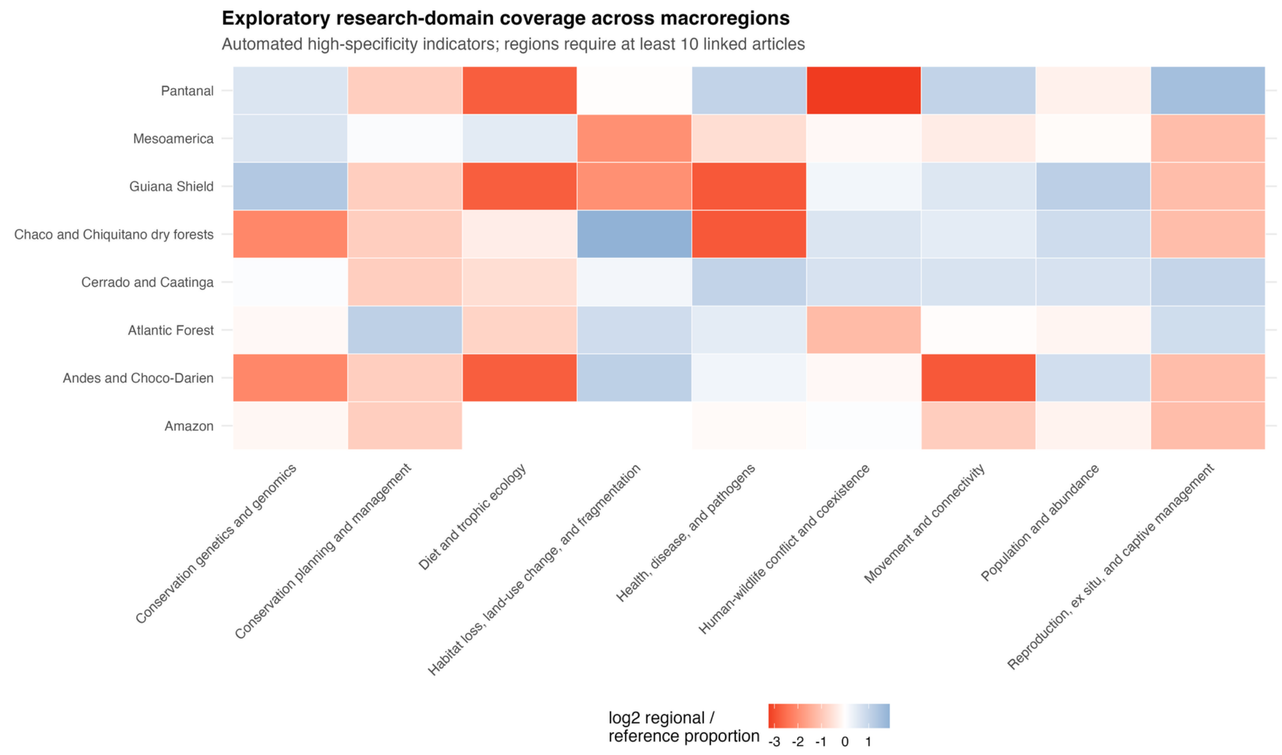


**Figure S10.** Exploratory relative coverage of metadata-derived research domains across broad regions. Cell values represent the log2 ratio between the regional domain proportion and the corresponding proportion in the complete georeferenced terrestrial reference set, after adding 0.01 to both proportions to avoid undefined values. Only regions represented by at least ten linked articles were included. This analysis was exploratory and did not contribute to candidate-gap classifications.

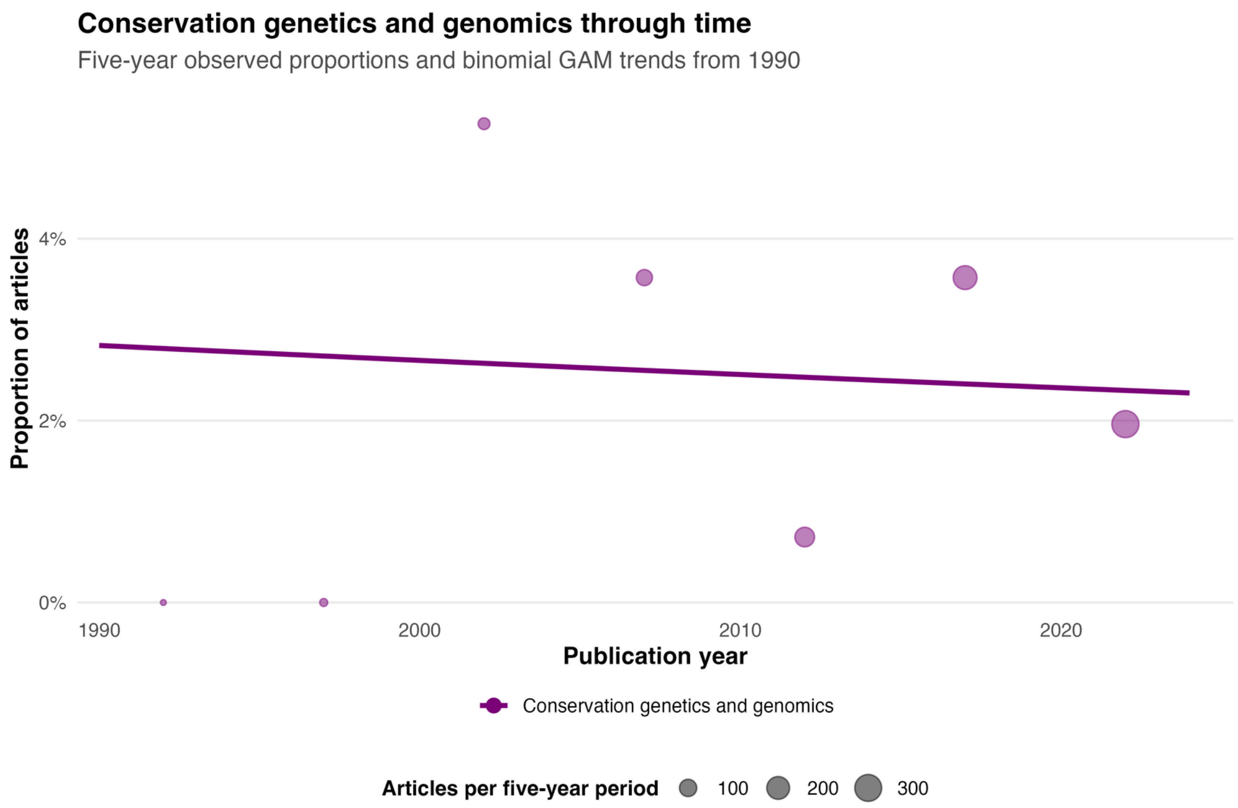


**Figure S11.** Temporal pattern of metadata-identified conservation genetics and genomics research. Points show observed proportions within five-year periods from 1990 onward, with point size proportional to the number of articles in each period. The line shows the fitted article-level binomial generalized additive model. The pattern is descriptive and should be interpreted cautiously because the domain contained only 20 identified articles.

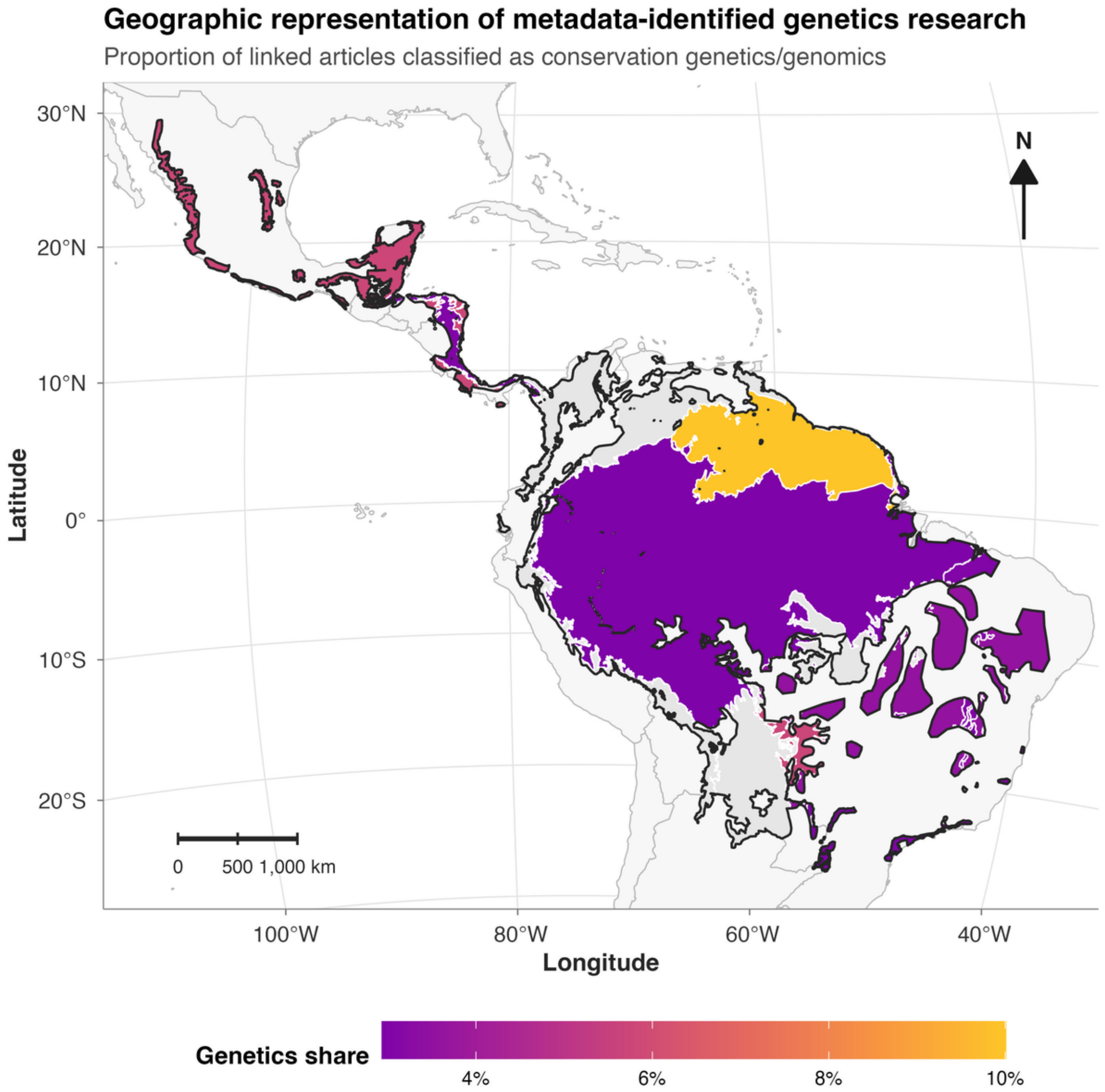


**Figure S12.** Geographic representation of metadata-identified conservation genetics and genomics research. Colors represent the proportion of linked articles in each broad region classified by the high-specificity conservation-genetics indicator; light gray indicates no linked genetics article. The map is exploratory because only eight of the 20 metadata-identified genetics articles were represented in the terrestrial spatial subset.

### SUPPLEMENTARY REFERENCES

References cited in Supplementary Material are included in the reference list of the main manuscript. The software citations listed in Table S4 should also be included in the main reference list when required by the target journal or retained within Table S4 when the journal permits software citations in supplementary tables.
